# AKT2 deficiency causes sarcopenia and metabolic disorder of skeletal muscle

**DOI:** 10.1101/805812

**Authors:** Miao Chen, Caoyu Ji, Fei Xiao, Dandan Chen, Shuya Gao, Qingchen Yang, Yue Peng, Daniel Sanchis, Fangrong Yan, Junmei Ye

## Abstract

Skeletal muscle is responsible for the majority of glucose disposal in the body. Insulin resistance in the skeletal muscle accounts for 85-90% of the impairment of total body glucose disposal in patients with tye 2 diabetes (T2D). However, the mechanism remains controversial. AKT2 is a protein kinase performing important functions in the regulation of glucose metabolism. We observed that mice deficient for AKT2 (*AKT2* KO) exhibited decreased body weight and lean mass and showed impaired glucose tolerance, compared to their age- and gender-matched wild type mice (WT). Therefore, to test whether *AKT2* deficiency causes deficits in skeletal muscle development and metabolism, we analyzed the expression of molecules related to skeletal muscle development, glucose uptake and metabolism in young (3 months) and old (8 months) mice. We found that AMPK phosphorylation and MEF2A expression were downregulated in young *AKT2* KO mice, and this downregulation was inverted by AMPK activation. We also observed reduced mtDNA abundance and reduced expression of genes involved in mitochondrial biogenesis in the skeletal muscle of adult *AKT2* KO mice, which was prevented by AMPK activation. However, GLUT4 expression was regulated by AKT2 in an AMPK-independent manner in skeletal muscle. During high-fat-diet (HFD)-induced obesity, *AKT2* KO mice exhibited increased insulin resistance compared to WT mice. Our study establishes a new and important function of AKT2 in regulating glucose uptake and AMPK-dependent development and mitochondrial biogenesis in skeletal muscle.

## Introduction

AKT, also known as protein kinase B (PKB) (1), is a serine/threonine kinase that plays a key role in insulin-stimulated glucose uptake (2). There are three isoforms of AKT family, AKT1/PKBα, AKT2/PKBβ and AKT3/PKBγ. The three AKTs are highly related to each other in both structure and amino acid sequence (∼85%) and are activated by phosphatidylinositol-3 kinase (3). The function of individual isoforms of AKT in skeletal muscle has been recognized by an increasing number of studies. Previous studies comparing the three proteins support the notion that these isoforms have similar biochemical characteristics (4). However, recent studies prove that the three isoforms of AKT exhibit tissue-specific expression and distinct functions. In mice, AKT1 loss of function causes diminished fetal growth but maintain normal glucose regulation (5). *AKT2* deficiency results in insulin resistance and a type 2 diabetes mellitus-like syndrome (6), and global knockout of AKT3 leads to reduced brain size (7). It is well documented that loss of *AKT2* is associated with metabolic disorder with glucose imbalance, growth deficiency (8, 9), loss of adipose tissue(10), but the mechanisms behind are not fully understood. Importantly, scarce knowledge is available regarding the alterations on regulatory mechanisms and molecular changes induced by *AKT2* deficiency in skeletal muscle, an active metabolic tissue that depends mainly on glucose as energy resource, and also a major site of insulin-stimulated glucose uptake.

After extensive proliferation, myoblasts undergo differentiation and fusion with each other or existing myofibers to build functional muscle tissue (11). Many transcription factors are involved in the control of myogenesis. Members of the MyoD family play important roles in the control of muscle differentiation during embryonic development and adult muscle regeneration (11). The myogenic activity of these transcriptional factors is enhanced through their interaction with the myocyte enhancer factor 2 (MEF2) family (11). In vertebrates, the MEF2 family includes four members, MEF2A, B, C and D, which share high sequence homology in their DNA binding and dimerization domains but are divergent in their C-terminal transcriptional activation domains (11). The MEF2 isoforms active in skeletal muscle are A, C and D (12). Skeletal muscle-specific deletion of MEF2C results in neonatal morality due to defects in muscle integrity and failure in sarcomere formation (13, 14). Moreover, combined deletion of the MEFA, C and D results in a blockade to generation, suggesting essential roles of MEF2A, C and D in satellite cell differentiation (11). It is reported that loss of *AKT2* results in growth deficiency by exhibiting decreased body weight (15–17), and impaired myotube maturation in the skeletal muscle (18). However, the impact of *AKT2* deficiency in muscle development and the possible implication of MEF2 transcription factors-dependent signaling has not been previously investigated.

Skeletal muscle is responsible for the majority of glucose disposal in the body. Impaired glucose handling capacity leads to a state of insulin resistance (19). Insulin resistance in the skeletal muscle accounts for 85-90% of the impairment of total body glucose disposal in patients with tye 2 diabetes (T2D) (20). *AKT2*-null mice exhibited insulin resistance with elevated plasma triglycerides (15) and AKT2 is essential for insulin-stimulated glucose uptake which cannot be replaced by AKT1 in mouse skeletal muscle (2). Interestingly, reduced AKT2 expression and impaired insulin-stimulated AKT2 activation are reported to occur in diabetic (21) and insulin resistant (22) human skeletal muscle. Thus, these researches suggest that AKT2 plays an important role in systemic glucose homeostasis.

AMP-activated protein kinase (AMPK) is a major regulator of energy substrate use in several organs. Responding to high AMP/ATP and ADP/ATP ratios, AMPK activation leads to enhanced fatty acid oxidation, thus influencing energy homeostasis (23). In addition, AMPK activation has been shown to promote the translocation of GLUT4 to the plasma membrane, thus stimulating glucose uptake in skeletal muscle (24–26). Furthermore, chronic activation of AMPK reduces markers of skeletal muscle fragility (27) and enhances muscle fiber oxidation capacity by stimulating mitochondrial biogenesis (28–30). It has been published that AKT1 negatively regulates phosphorylation of AMPK in the myocardium and MEFs (31, 32). Additionally, it is reported that MEF2A is activated in response to AMPK activation in skeletal muscle (33). However, whether AKT2 functions as regulator of AMPK and the regulatory signaling pathway in normoxic skeletal muscle remains to be investigated.

We observed that *AKT2* KO mice exhibited decreased body weight compared to their age- and gender-matched WT mice. Despite obvious loss of white adipose tissue as previously reported (2, 15, 34), we also found that the lean mass of *AKT2* KO mice decreased significantly. These data indicate an important role for AKT2 in skeletal muscle development. Therefore, we hypothesized that *AKT2* deficiency causes deficits in skeletal muscle development, which leads to the alteration of skeletal muscle reprogramming in development and growth. To test this, we analyzed the expression of molecules related to skeletal muscle development, glucose uptake and metabolism at both young (3 months) and old (8 months) mice. We found that upon AMPK activity and MEF2A were downregulated in young *AKT2* KO mice, and this downregulation was prevented by AMPK activation. We also observed that mitochondrial biogenesis and oxidative capacity were impaired in the skeletal muscle of *AKT2* deficient mice. In addition, *AKT2* KO mice exhibited much deteriorated insulin resistance after high-fat-diet (HFD) feeding. Our results indicate that AKT2 is an important factor for AMPK activation and mitochondrial biogenesis in the skeletal muscle, and thus establish a new important function of AKT2 in the development and glucose metabolism of skeletal muscle.

## Results

### 1. Disturbance of glucose metabolism due to AKT2 deficiency

To investigate the impact of *AKT2* deficiency on growth, body mass of mice at different ages was recorded. *AKT2* KO mice were born healthy, and were able to grow to adulthood without abnormalities of lifespan, as previously described by other groups (35). However, *AKT2* KO mice showed a reduction in body weight beginning around postnatal day 40 (Fig. 1A), and from P90, the *AKT2* KO mice were visibly distinguishable from their WT littermates (Fig. 1B), although there were no differences of tibial length between WT and *AKT2* KO mice (suppl. Fig.1), nor in the daily food intake (Fig. 1C and D).

**Figure 1.**
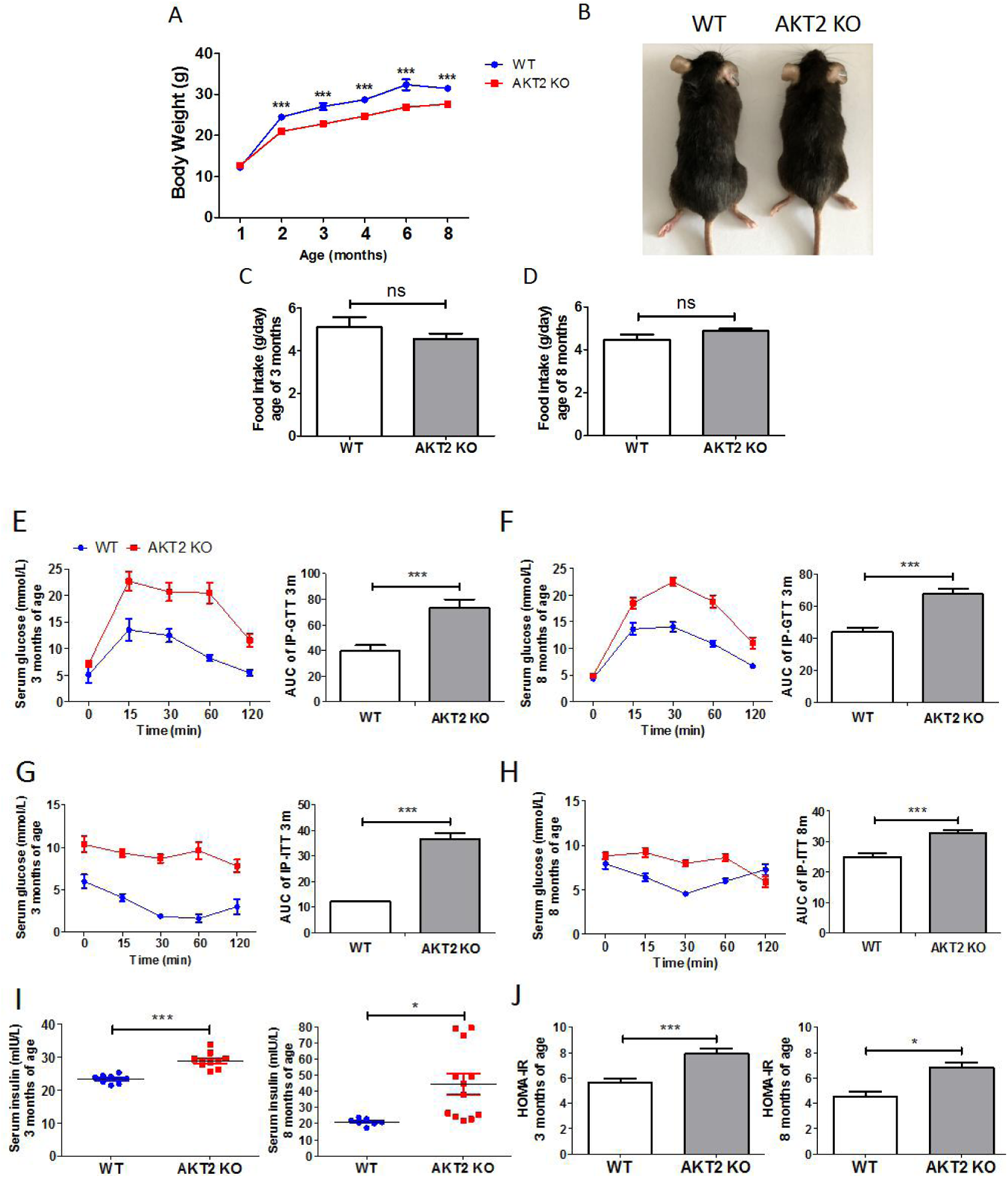
Impaired glucose metabolism of *AKT2* KO mice. (**A**) Body weight curve of WT and *AKT2* KO mice at different ages that range from 1 to 8 months old (WT, n=10; *AKT2* KO, n=15). (**B**) Representative images of *AKT2* KO mice with WT littermates at the age of 8-month-old. (**C and D**) Food intake of mice at the age of 3- and 8-month-old (n=10 for both genotypes). (**E and F**) Glucose tolerance with AUC of both genotypes at the age of 3-month-old (WT, n=6; *AKT2* KO, n=11) and 8-month-old (n=10 for both genotypes). (**G and H**) Insulin tolerance with AUC of both genotypes at the age of 3-month-old (n=5 for both genotypes) and 8-month-old (n=4 for both genotypes). (**I and J**) Serum insulin and HOMA of insulin resistance of both genotypes at the age of 3-month-old (WT, n=8; *AKT2* KO, n=11) and 8-month-old (WT, n=7; *AKT2* KO, n=9). \**P* < 0.05; \*\*\**P* < 0.001, WT versus KO. Statistical significance was determined using unpaired 2-tailed Student’s *t* test.

Since AKT2 functions downstream of PI3K which mediates insulin-induced effects on glucose metabolism (6, 15, 36), we investigated whether glucose metabolism was also affected due to *AKT2* deficiency by evaluating global glucose metabolic status. *AKT2* KO mice exhibited impaired glucose tolerance at 3 and 8 months of age (Fig. 1E and F; suppl. Fig. 2). Interestingly, glucose intolerance was less pronounced in female *AKT2* KO mice compared to age-matched male mice (suppl. Fig.2). Insulin sensitivity was also impaired in *AKT2* KO at 3 and 8 months of age (Fig. 1G and H). Plasma insulin levels and HOMA-IR were elevated in *AKT2* KO mice at 3 and 8 months of age (Fig. 1I and J). These results suggest that *AKT2* deficiency in skeletal muscle impairs both the response of glucose tolerance and insulin sensitivity.

### 2. Tissue and age-related changes in AKT2 expression

Next we explored AKT2 expression in different tissues. AKT2 mRNA and protein were enriched in tissues highly associated with glucose metabolism including pancreas, liver, brain and soleus muscle (Fig. 2A and B). To further investigate the expression pattern of AKT2 in skeletal muscle during development, AKT2 protein expression in soleus muscle of mice at different ages was evaluated. We observed a positive correlation between p-AKT2 expression and age in this tissue (Fig. 2C). To verify this result, we also checked p-AKT2 and AKT2 in C2C12 myofibroblasts at different differentiating days. In agreement with the result in vivo, p-AKT2 increased during the differentiation of C2C12 cells (Fig. 2D), suggesting that AKT2 activity plays an important role in the development of skeletal muscle. To determine if there is compensatory effect of AKT1 due to AKT2 absence, we assessed the abundance of AKT1 and AKT2 in soleus muscle of both WT and *AKT2* KO mice. The result showed that AKT1 expression was similar in skeletal muscle of both genotypes (Fig. 2E), suggesting that there was no compensatory expression of AKT1 in the skeletal muscle deficient for *AKT2*.

**Figure 2.**
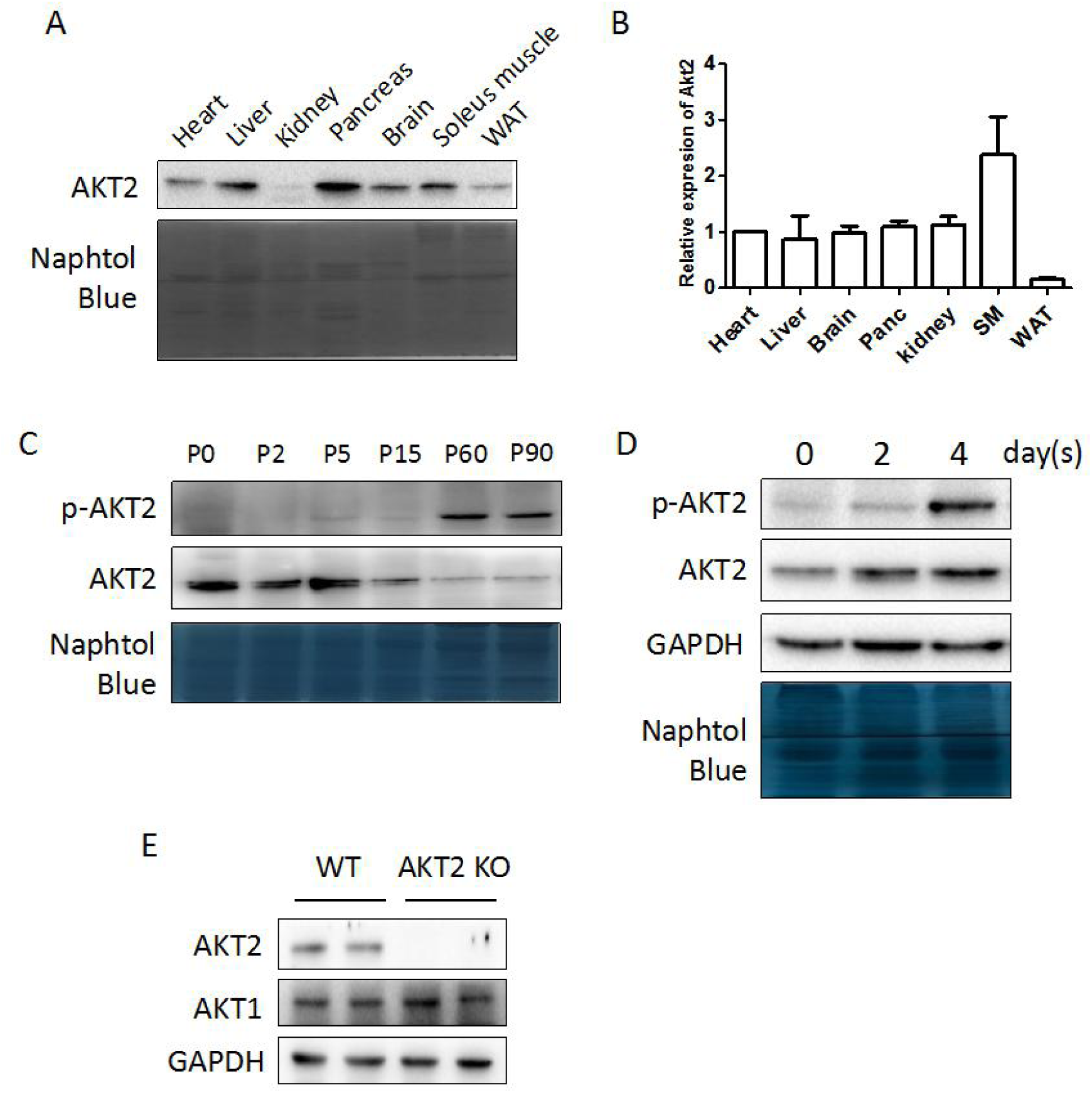
AKT2 expression in different tissues and ages. (**A and B**) Western blot and real-time PCR analysis of AKT2 protein abundance in multitissue panel using a C57BL/6 mouse. (**C**) p-AKT2 and AKT2 protein expression in skeletal muscle from embryo throughout development and into adulthood (n=3). (**D**) p-AKT2 and AKT2 protein expression in C2C12 cells at different days during differentiation. Representative immunobolts are shown from n=3. (**E**) AKT1 and AKT2 expression in skeletal muscle of WT and AKT2 KO mice at the age of 3-month-old (n=4 for both genotypes). Statistical significance was determined using unpaired 2-tailed Student’s *t* test.

### 3. Loss of skeletal muscle mass of mice deficient for AKT2

The analysis of tissue weight of WT and *AKT2* KO mice at 3 and 8 months of age confirmed the reduction of WAT mass, as previously described (2,12,31), yet demonstrated also a reduction of lean mass of *AKT2* KO mice, compared to WT (Fig. 3 A; Suppl. Figure 3A, B and C), without obvious morphological changes of pancreas at 8 months of age (Suppl. Figure 4). To assess the role of AKT2 in skeletal muscle growth, we weighed different muscle groups in 3-month-old and 8-month-old mice. The hind limb soleus, quadriceps, gluteus and gastrocnemius, all weighed significantly less in *AKT2* KO mice compared to age-matched WT mice (Fig. 3B and C). Further histological analysis revealed normal nuclear position and similar H&E staining of either longitudinal or transverse sections (Fig. 3D). However, the cross-sectional area of soleus muscle from *AKT2* KO mice was significantly reduced compared to WT (Fig. 3E), and this change in the mean cross-sectional area of myofibers coincided with a decrease in skeletal muscle mass of 3-month-old *AKT2* KO mice. Electron microscopy analysis also revealed smaller sarcomeres in the muscle of *AKT2* KO mice (Fig. 3F), suggesting impaired myotube development. However, there were no abnormalities in Z-discs and myofibrils (Fig. 3F). These results suggested that, in *AKT2-*deficient mice, the reduction of body weight, which coincides with a decrease in the lean mass at the age of 3 months and 8 months, was partly caused by loss of muscle weight, suggesting an important role for AKT2 in regulating skeletal muscle development.

**Figure 3.**
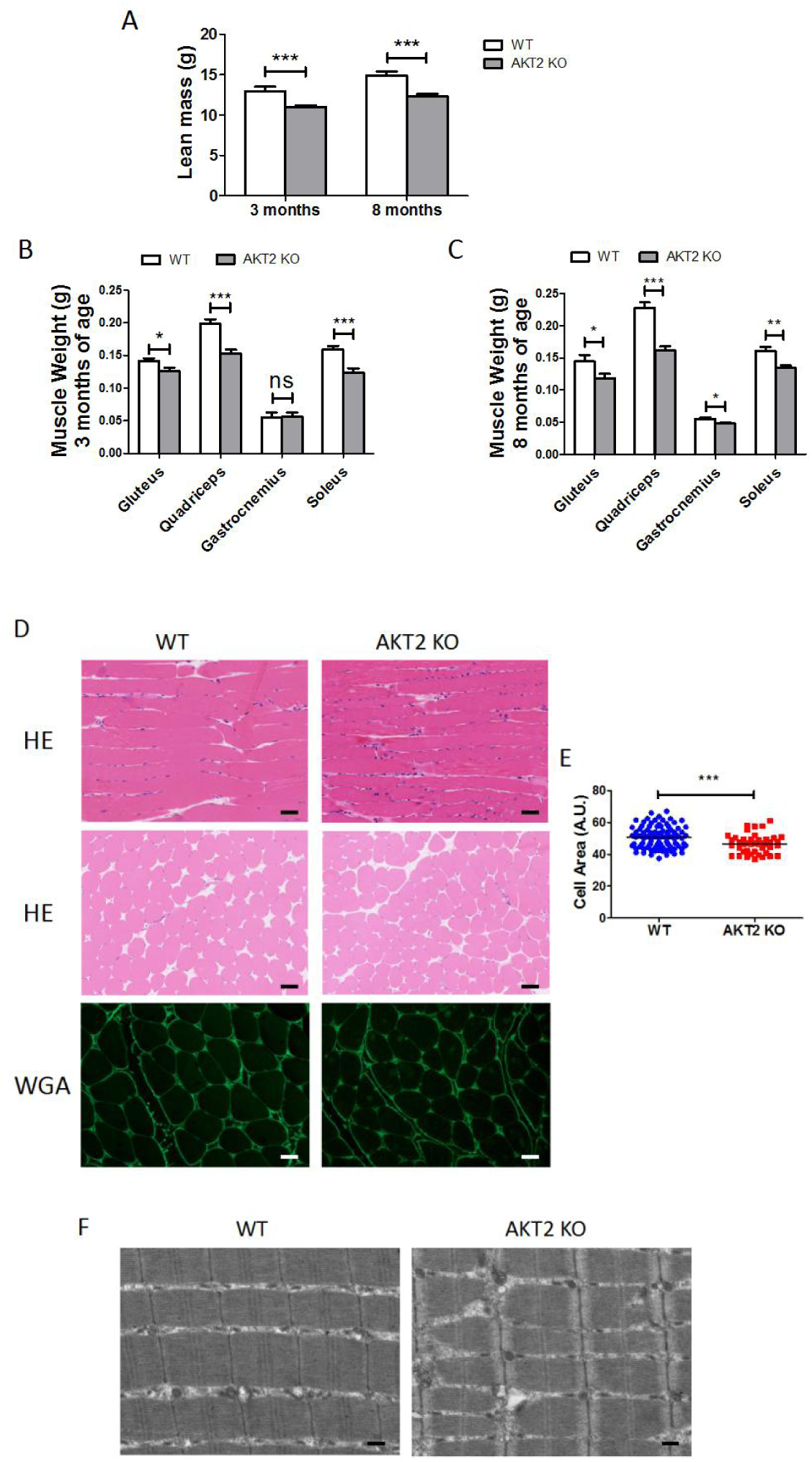
Sarcopenia of AKT2 KO mice. (**A**) Lean mass of mice at the age of 3- (n=15 for both genotypes) and 8-month-old (n=12 for both genotypes). (**B and C**) Muscle weight of different parts from mice hindlimb at the age of 3- (WT, n=10; *AKT2* KO, n=8) and 8-month-old (n=10 for both genotypes). (**D**) H&E staining of Longitudinal (upper panel) and transverse sections of soleus muscle (middle panel), and WGA staining of soleus muscle (lower panel) from both genotypes (WT, n=4; *AKT2* KO, n=8). scale bar: 100 μm for H&E; 50 μm for WGA. (**E**) Muscle area calculation based on WGA staining of soleus muscle (WT, n=4; *AKT2* KO, n=6). (**F**) Representative transmission electron microscopic images showing regular sarcomeric morphology in WT and *AKT2* KO soleus muscle (n=3 per genotype). scale bar: 4 μm \**P* < 0.05; \*\**P* < 0.01; \*\*\**P* < 0.001, WT versus KO. Statistical significance was determined using unpaired 2-tailed Student’s *t* test.

### 4. Retardation of Skeletal muscle development due to AKT2 loss-of-function

AMPK serves as an important regulator to increase muscle development (37). In addition, AMPK and MEF2 are shown to cooperate in the transcriptional regulation of genes related to glucose uptake (33). We next sought to determine whether muscular AMPK/MEF2 signaling was altered in *AKT2* KO mice. The phosphorylation of AMPK (p-AMPK) and the expression of MEF2A were significantly decreased in *AKT2* KO muscle (Fig. 4A and B). Further investigation of the above events in the C2C12 myoblast cell line showed that, as expected, MEF2A expression increased during *in vitro* differentiation (Fig. 4C). To investigate the existence of a direct link between *AKT2* deficiency and AMPK hypophosphorylation, C2C12 myoblasts were transfected with either lentivirus carrying empty vector (scr) or lentivirus carrying small interference RNA knockdown of *AKT2* (si*AKT2*). The results showed that p-AMPK decreased in cells transfected with siA*KT2* compared to controls at day 7 post differentiation (Fig. 4D). In agreement with the *in vivo* observations, MEF2A was less abundant in si*AKT2* treated cells (Fig. 4D). Moreover, morphological analysis of C2C12 cells also suggested that *AKT2* knockdown resulted in retardation of cell differentiation (Fig. 4E). These results indicate that the AMPK-MEF2A axis is inhibited in skeletal muscle due to *AKT2* loss-of-function.

**Figure 4.**
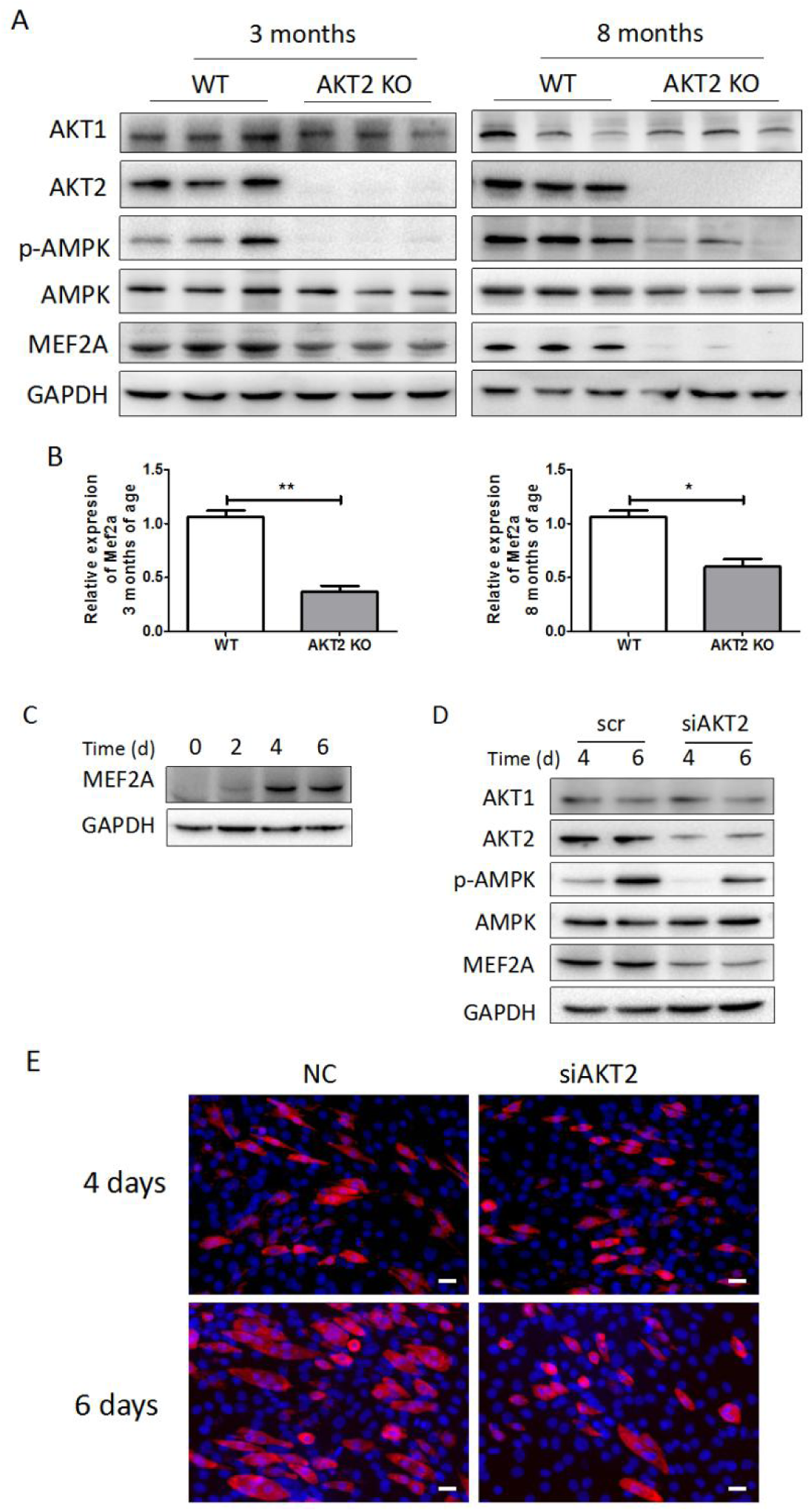
Suppression of AMPK-MEF2A signaling due to AKT2 deficiency. (**A**) Representative immunoblots of AKT1, AKT2, p-AMPK, AMPK, MEF2A in soleus muscle of mice from both genotypes at the age of 3- and 8-month-old (n=8 for both genotypes). (**B**) Relative mRNA expression of mef2a in skeletal muscle (n=5 for both genotypes). (**C**) Representative immunoblots of AKT1, AKT2 and MEF2A in C2C12 cells at different differentiating days. (**D**) Representative immunoblots of AKT1, AKT2, p-AMPK, AMPK and MEF2A in C2C12 cells transfected with either negative control (NC) or siAKT2. (n=4). (**E**) Representative immunofluorescence staining of C2C12 cells with α-actinin and DAPI at day 4 and day 6 after transfection with either NC or siAKT2 (n=4) (Blue: nucleus; Red: α-actinin). scale bar: 20 μm. Statistical significance was determined using unpaired 2-tailed Student’s *t* test.

### 5. Alterations of mitochondrial biogenesis and function in skeletal muscle of AKT2 KO mice

The capacity of the muscle to import and utilize energy is an important determinant of its function, and mitochondria play a central role in oxidative metabolism. The high levels of AKT2 expression in metabolically active organs and the link with AMPK signaling pointed to a previously unappreciated effect of AKT2 in mitochondrial function and metabolism. Therefore, we sought to determine whether relevant mitochondrial parameters were altered in skeletal muscle deficient for *AKT2*. Electron microscopy revealed no gross morphological changes of mitochondria (suppl. Fig. 5). We also quantified mitochondrial DNA (mtDNA)/genomic DNA, and the result showed diminished mtDNA copies in the 8-month-old *AKT2* KO mouse muscle (Fig. 5A and B). Peroxisome proliferator activated receptor gamma coactivator 1 alpha (PGC-1α), mitochondrial transcription factor A (MTFA), NRF1 and Endonuclease G (EndoG) are recognized as regulators of mitochondrial function and biogenesis (38–42). Our results showed that *AKT2* deficiency results in decreased expression of *pgc-1α*, m*tfa*, *nrf-1* and *pparα* in the skeletal muscle deficient for *AKT2* both at the transcript (Fig. 5C) and protein (Fig. 5D) levels, suggesting that impaired mitochondrial biogenesis and function due to *AKT2* deficiency.

**Figure 5.**
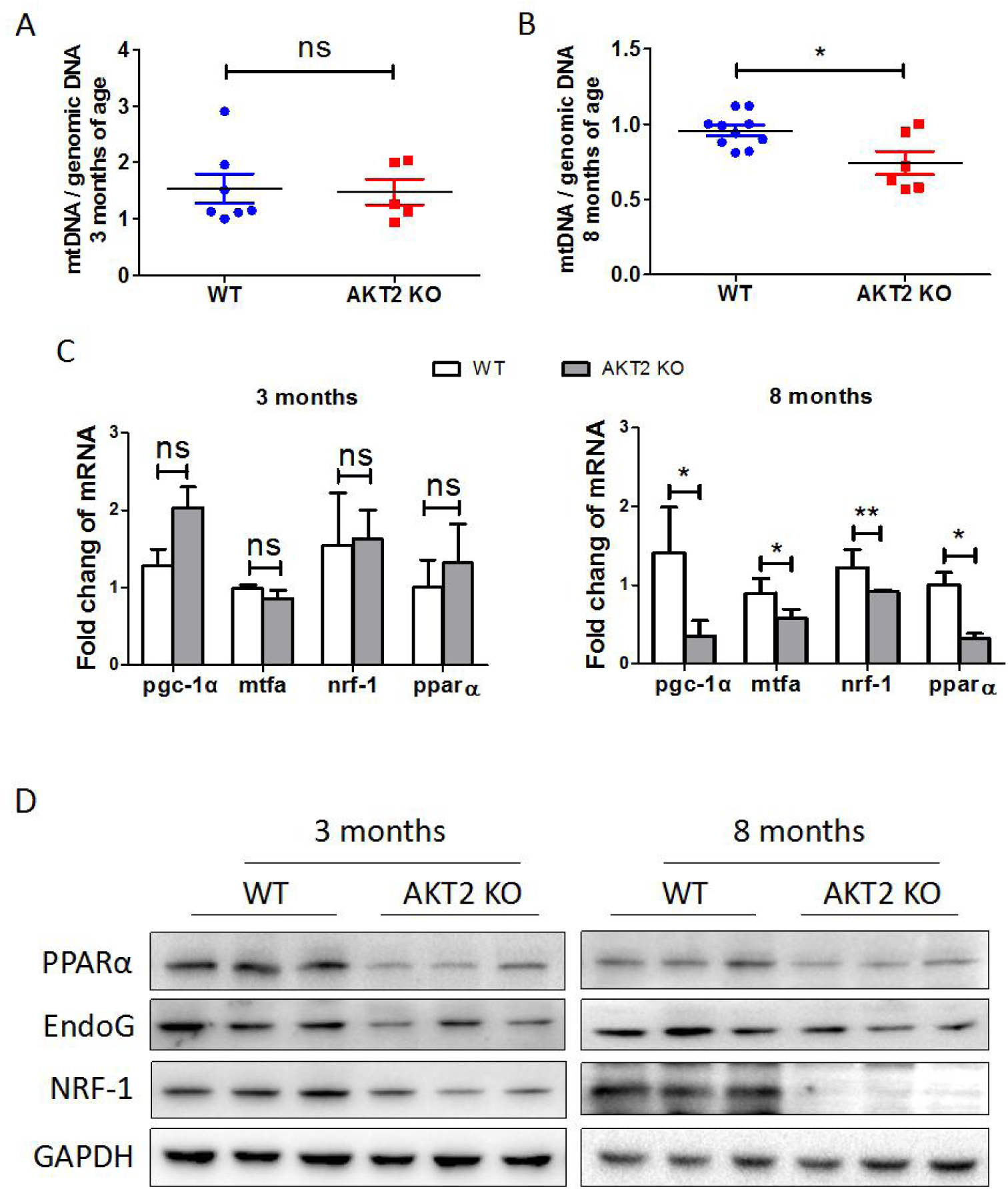
Impaired mitochondrial biogenesis of skeletal muscle with AKT2 deficiency. (**A and B**) Quantification of mitochondrial copies in the skeletal muscle of WT and AKT2 KO mice at the age of 3- (WT, n=7; *AKT2* KO, n=5) and 8-month-old (WT, n=10; *AKT2* KO, n=6). The abundance of mitochondrial DNA was measured by real-time PCR to determine the copy number shown as the ratio of *nd-5* vs *β-actin*. (**C**) Expression of mRNA for *pparα*, *pgc-1α*, *mtfa* and *nrf1* in the skeletal muscle tissue of mice at the age of 3- and 8-month-old (n=5 for both genotypes). (**D**) Representative immunoblots of PPARα, EndoG and NRF-1 in soleus muscle from both genotypes at the age of 3- and 8-month-old (n=8 for both genotypes). \**P* < 0.05; \*\**P* < 0.01; ns, not significant, WT versus KO. Statistical significance was determined using unpaired 2-tailed Student’s *t* test.

### 6. Disturbance of glucose uptake and usage in skeletal muscle of AKT2 KO mice

In healthy subjects, skeletal muscle is responsible for consuming nearly 80% of insulin-stimulated glucose uptake (43), suggesting that the skeletal muscle is a major regulator of glucose metabolism. In skeletal muscle, the insulin-stimulated glucose uptake is performed by GLUT4, which rapidly translocated to the plasma membrane in response to insulin stimulation (44). Since *AKT2* KO mice exhibited hyperglycemia at an early stage of life, we decided to explore whether there were defects on glucose uptake in the muscle and analyzed GLUT4 expression in this tissue. Our result showed that GLUT4 protein abundance was significantly decreased in soleus muscle of *AKT2* KO mice at the age of 3 and 8 months (Fig. 6A, C, D and F). We next evaluated the activity of glycogen synthase kinase 3 beta (GSK3β), a serine/threonine kinase essential for the regulation of glycogen synthesis (45). Our result showed that, in addition to markedly reduced GLUT4 expression, GSK3β phosphorylation (p-GSK3β) was increased by 30% in 3-month-old *AKT2* KO skeletal muscle compared to WT (Fig. 6A, B, D and E). Interestingly, at the age of 8 months, p-GSK3β in skeletal muscle of *AKT2* KO mice increased as much as 20% compared to WT (Fig. 6B and E). To verify the regulation of GLUT4 by AKT2, GLUT4 was checked in C2C12 cells transfected with NC or si*AKT2*. The result showed that GLUT4 expression decreased at both protein and mRNA level after *AKT2* knockdown (Fig. 6 G-I), suggesting that AKT2 positively regulates GLUT4 expression in skeletal muscle.

**Figure 6.**
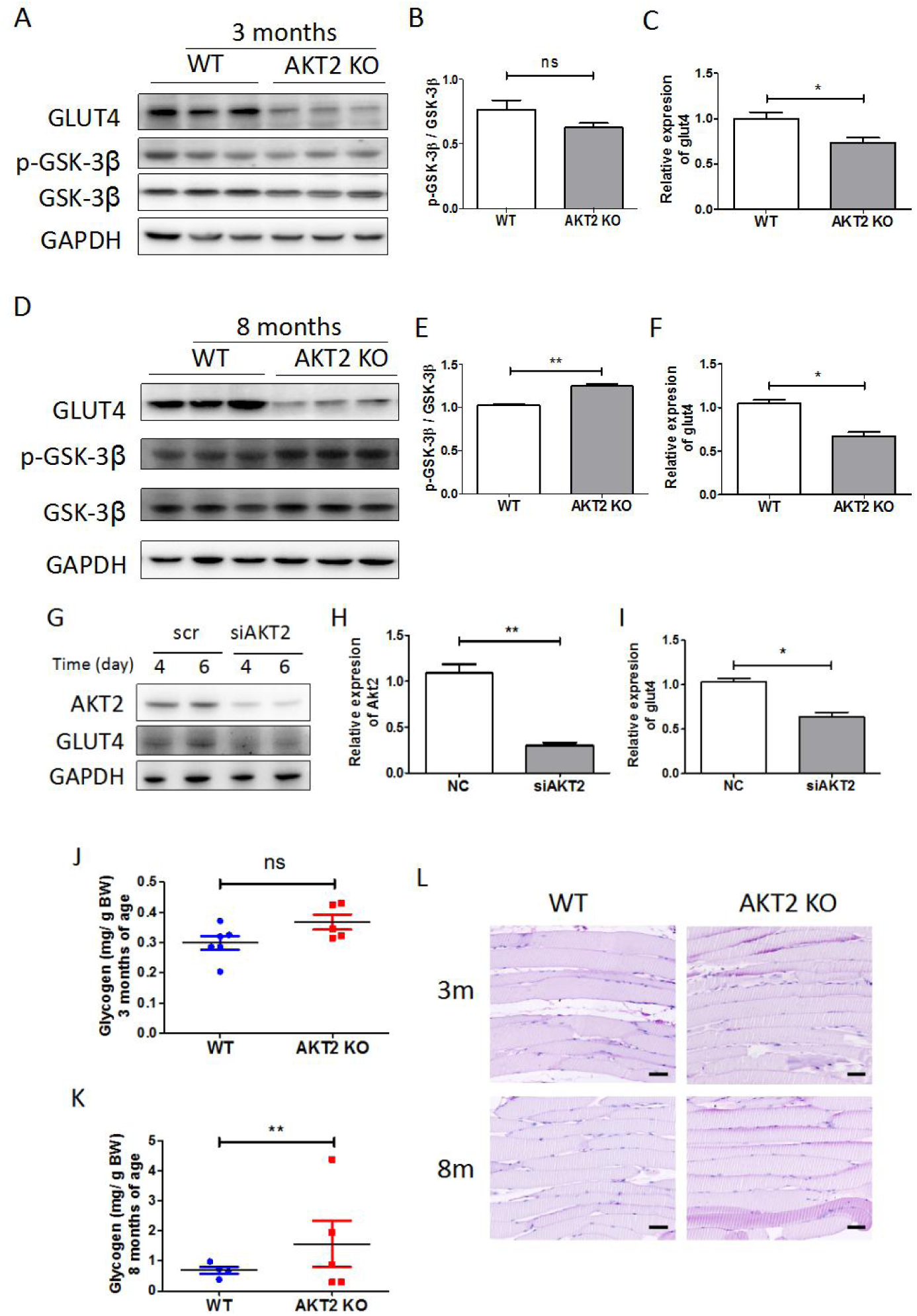
Glucose metabolic disorder in skeletal muscle with AKT2 deficiency. (**A-F**) Representative immunoblots of GLUT4, p-GSK3β and GSK3β in soleus muscle from WT and *AKT2* KO mice, and calculation of p-GSK3β/GSK3β ratio at the age of 3-month-old and 8-month-old (n=6 for both genotypes). (**G**) Representative immunoblots of AKT2, GLUT4 in C2C12 cells at different days after si*AKT2* transfection (n=3). (**H and I**) Real-time PCR of *akt2* and *glut4* in C2C12 cells at 6 days after si*AKT2* transfection (n=3). (**J and K**) Quantification of glycogen content in soleus muscle of mice from both genotypes at the age of 3- (WT, n=6; *AKT2* KO, n=5) and 8-month-old (WT, n=4; *AKT2* KO, n=5). (**L**) Glycogen distribution shown by PAS staining in soleus muscle of mice from WT and AKT2 KO mice at the age of 3- and 8-month-old (n=4 for both genotypes). scale bar: 100 μm. \*\**P* < 0.01; ns, not significant, WT versus KO. Statistical significance was determined using unpaired 2-tailed Student’s *t* test.

To check the amount of glycogen stored in skeletal muscle of *AKT2* KO mice, glycogen quantification and staining were performed in both genotypes of 3- and 8-month old mice. The results showed that the glycogen quantity was identical in soleus muscle of WT and *AKT2* KO younger mice, but there was an increase of glycogen content in the skeletal muscle of *AKT2*-deficient 8-month-old mice (Fig. 6J-L). These data suggest that, on one hand *AKT2* deficiency accounts for decreased glucose uptake and disturbance of carbohydrate metabolism in skeletal muscle, which may be a main cause of increased blood glucose. On the other hand, increased expression of p-GSK3β and glycogen content in AKT2 KO adult but not young mice suggest that altered glycogen metabolism is a secondary effect due to *AKT2* loss-of-function.

### 7. Improvement of global and skeletal muscle metabolism by AMPK activator in AKT2-null mice

To check the potential role of AMPK in *AKT2* deficiency-induced skeletal muscle sarcopenia, changes in glucose uptake, insulin resistance and mitochondrial biogenesis, an AMPK activator, AICAR, was continuously injected in *AKT2* KO or WT mice for 4 weeks. Glucose and insulin tolerance assays were performed at the end of AICAR treatment. The results showed that *AKT2* KO mice treated with AICAR had significantly improved insulin tolerance compared to untreated mice (Fig. 7A and B), without affecting body weight (Fig. 7C). Furthermore, as an indication of AMPK activity, we measured phospho-ACC (p-ACC) levels in soleus muscle and found that p-ACC increased two-fold post-AICAR treatment (Fig. 7D). To investigate the possible activation of the MEF2 signaling pathway by AMPK, we assessed the expression of MEF2A, MEF2D and factors related to glucose metabolism and mitochondrial biogenesis in this setting. The results showed that expression of MEF2A was significantly induced by AICAR treatment compared to saline-treated controls (Fig. 7D). Moreover, phosphorylation of GSK3β, which was highly stimulated in *AKT2*-deficient samples, was normalized by AICAR infusion (Fig. 7D). In addition, there was also significant elevation of NRF-1, PPARα, and EndoG protein abundance (Fig. 7D). Therefore, our data suggest that the decrease of MEF2A, as well as molecules involved in glucose metabolism and mitochondrial biogensis in skeletal muscle deficient for *AKT2* is due to AMPK inactivation, which may in turn contribute to aggravate insulin resistance and to impair glucose tolerance in serum.

**Figure 7.**
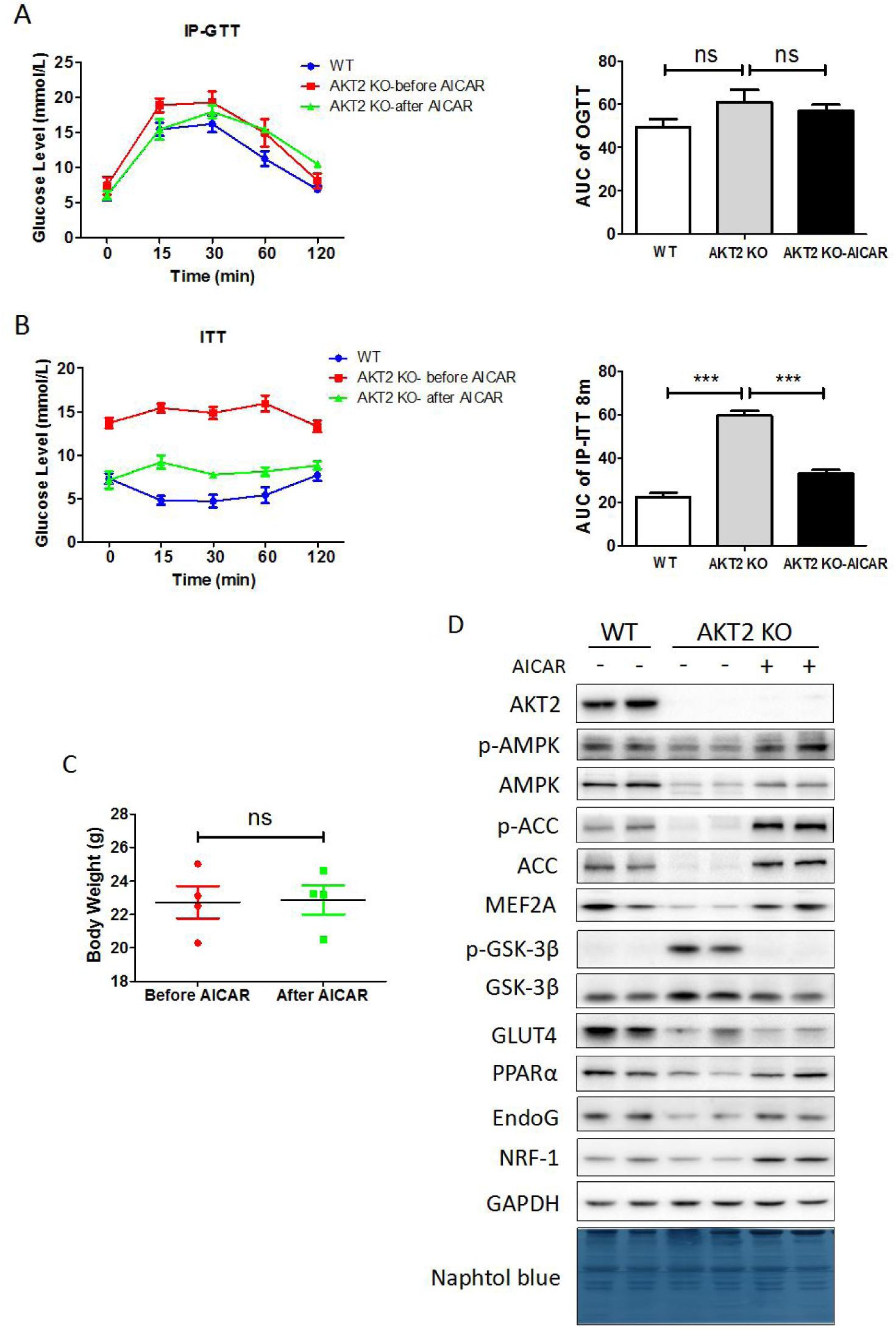
Reverse of impaired skeletal muscle develop and glucose metabolism by AICAR in AKT2 KO mice. (**A**) Glucose tolerance were evaluated before and after 4 weeks’ AICAR application to AKT2 KO mice and calculation of AUC (n=4 for both treatment). (**B**) Insulin tolerance were evaluated before and after 4 weeks’ AICAR application to AKT2 KO mice and calculation of AUC (n=4 for both treatment). (**C**) Body weight of AKT2 KO mice before and after AICAR application (n=4 for both treatment). (**D**) Representative immunoblots of soleus muscle from 6-month-old WT, AKT2 KO and AKT2 KO treated with AICAR mice (n=4 for both treatment). \*\*\**P* < 0.001; ns, not significant. Statistical significance was determined using unpaired 2-tailed Student’s *t* test.

### 8. Upregulation of AKT2 activity in skeletal muscle with insulin resistance

To investigate the function of AKT2 on global metabolic homeostasis, WT and *AKT2* KO mice were fed high fat diet (HFD; see methods for composition), for 12 weeks. At the end of this period, mice of the four groups were submitted to glucose tolerance test and insulin tolerance test. *AKT2* KO mice with regular chow exhibited insulin resistance and decreased glucose tolerance (Fig. 8A and B). HFD feeding induced impaired glucose tolerance in both genotypes at almost the same extent (Fig. 8A and B). However, although WT mice showed insulin resistance after HFD feeding for 12 weeks, HFD induces much greater impairment of insulin resistance in *AKT2* KO mice (Fig. 8C and D). Serum insulin was also highly induced in *AKT2* KO mice with HFD treatment (Fig. 8E and F). Therefore, our results suggest that AKT2 plays a key role in maintaining metabolic homeostasis of glucose and proper insulin response during HFD-induced obesity. We then compared the expression of p-AKT2, AKT2, p-AMPK and AMPK in the soleus muscle of WT mice fed normal diet (Vehicle) or HFD. The result showed that compared to the skeletal muscle from Vehicle group, the skeletal muscle of HFD-treated mice exhibited increased p-AKT2 protein and decreased total AKT2 protein abundance, and p-AMPK decreased in soleus muscle of mice with treated HFD (Fig. 8G). These data indicate that during the development of obesity and insulin resistance caused by diet, increased p-AKT2/AKT2 ratio may be accused for glucose metabolic disorder in skeletal muscle.

**Figure 8.**
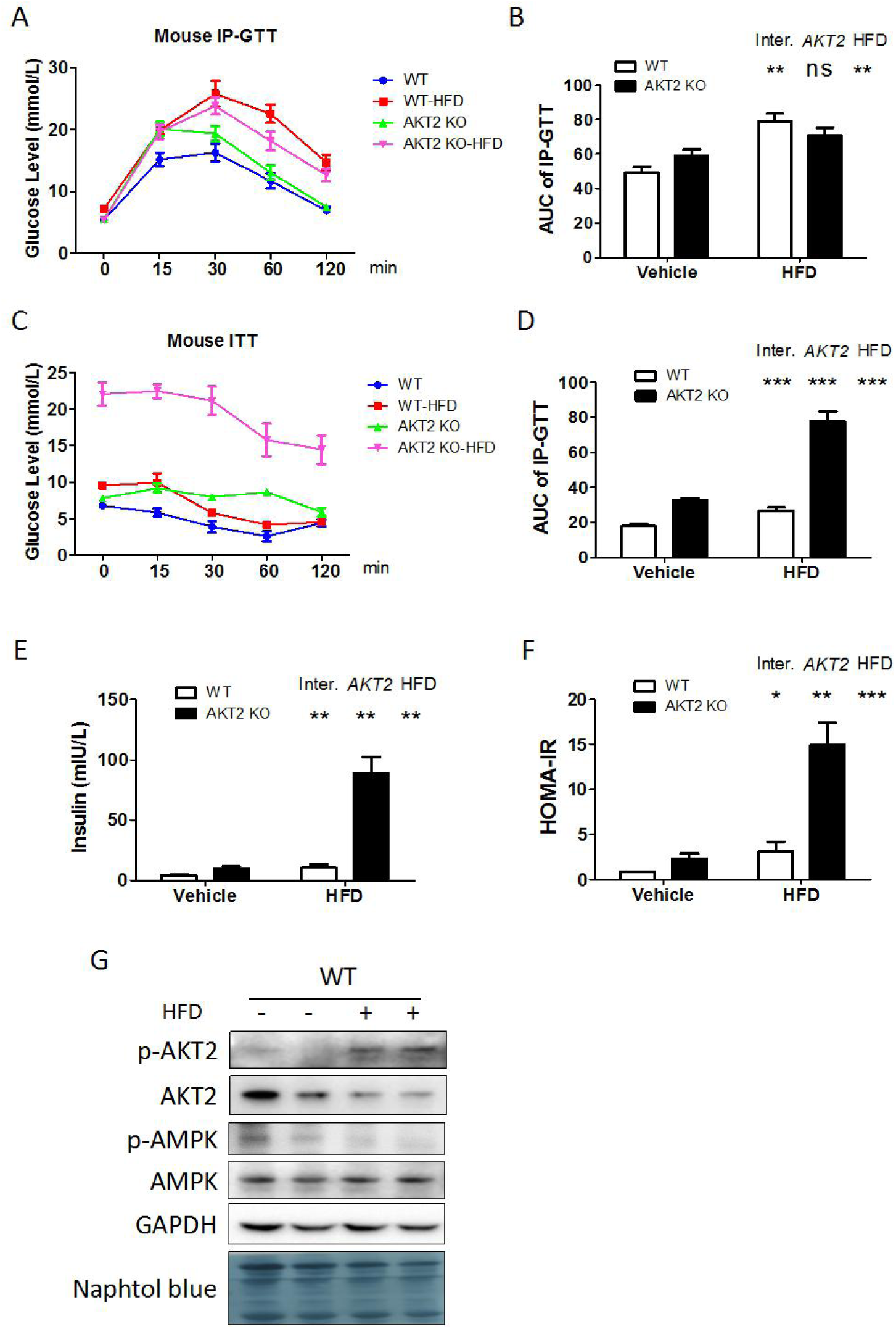
Decrease of AKT2 abundance in skeletal muscle of mice with insulin resistance. (**A-D**) WT and *AKT2* KO mice were fed ND or HFD for 12 weeks, glucose tolerance and insulin tolerance were evaluated at the end of 12-week feeding with ND or HFD (n=6 for both treatment). (**E**) Serum insulin concentration of mice with different treatment and (**F**) HOMA-IR calculation (n=4-9). (**G**) Representative immunoblots of p-AKT2, AKT2, p-AMPK and AMPK in skeletal muscle from mice fed ND or HFD for 12 weeks (n=4 for both treatment). \**P* < 0.05; \*\**P* < 0.01; \*\*\**P* < 0.001, WT versus KO. Statistical significance was determined using two-way ANOVA followed by Bonferroni post-test.

**Figure 9.**
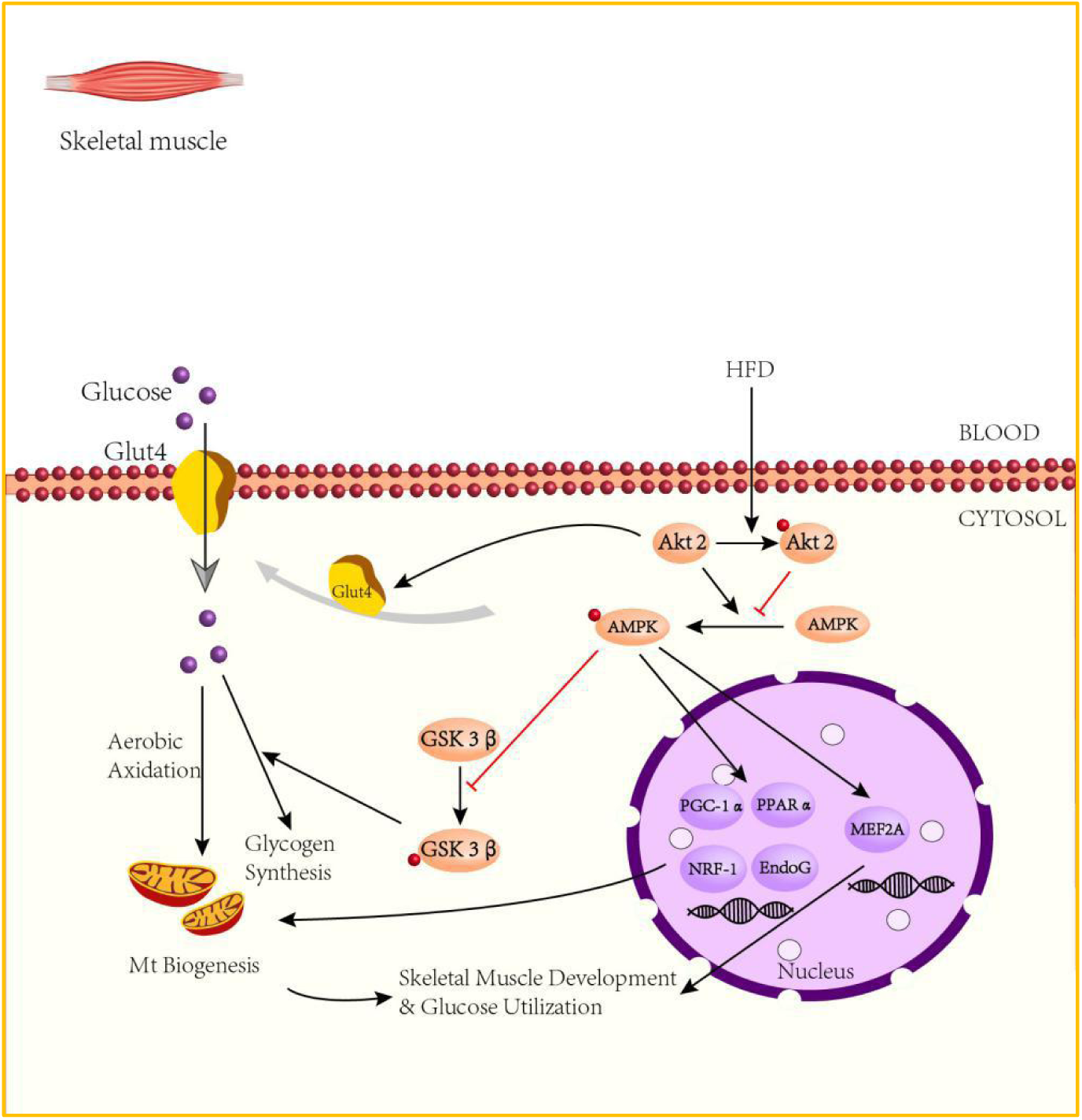
Schematic of AKT2 at a skeletal muscle in the network of gene regulation that underpins the program of muscle development and metabolism. During muscle development, AKT2 promotes GLUT4 expression, which transports glucose from blood into muscle cells. AKT2 also increases AMPK phosphorylation, which promotes the expression of MEF2A that positively regulate skeletal muscle development. However, when AKT2 is phosphorylated by stimulations such as high-fat diet (HFD), it has the opposite function. On the other hand, activated AMPK promotes expression of genes related to mitochondrial biogenesis, including PPARα, PGC-1α, NRF-1 and EndoG. p-AMPK also inhibits phosphorylation of GSK3β upon AKT2 regulation, therefore increases glycogen synthesis in skeletal muscle cells.

## Discussion

Multiple lines of evidence support the hypothesis that AKT2 is involved in growth and glucose metabolism (2, 6, 46). In this study, we provide evidence that AKT2 positively regulates AMPK activation, which play an important role in the control of growth, mitochondrial biogenesis and function of skeletal muscle. We found that p-AMPK, MEF2A, and molecules associated with mitochondrial biogenesis and function were downregulated in skeletal muscle of *AKT2* KO mice, which can be reversed by AMPK activator. We also observed that AKT2 regulates GLUT4 expression in an AMPK-independent manner in skeletal muscle. Moreover, disturbed glucose metabolism was more severe in *AKT2* KO-HFD mice, and activation of AKT2 activity was observed in WT mice with HFD-induced obesity. Our results indicate that AKT2 is involved in AMPK activation which in turn regulates skeletal muscle development. We also proved that AKT2 participates in mitochondrial biogenesis and glucose metabolism at normoxic skeletal muscle. Therefore, our study establishes a new and important role for AKT2 in the developmental and metabolic regulation of glucose in skeletal muscle.

*AKT1* and *AKT2* KO mice exhibit remarkably different phenotypes (5, 47–49), suggesting that the two isoforms regulate a subset of unique downstream genes and biological functions respectively. First, we checked AKT2 expression in different tissues of mice, the results showed that AKT2 is most abundant in organs that are active for glucose metabolism. We further found elevated p-AKT2 protein expression in soleus muscle at different ages. This result was also verified *in vitro* by checking different days of differentiation in C2C12 cells, indicating important role of AKT2 during skeletal muscle development. In *AKT2* KO mice, there was no difference of AKT1 protein abundance compared to WT, suggesting the unique function of AKT2 in skeletal muscle, which is not compensated by other AKT isoforms.

In *AKT2* KO mice, we observed constant mild decrease of body weight since the age of 1-month-old and throughout the life, which is in agreement with other research (15, 17). Since most research reported that the loss of body weight in *AKT2* KO mice is largely due to decreased white adipose tissue (WAT), few focused on the alteration of skeletal muscle. Therefore, to investigate whether there was impact of *AKT2* loss-of-function on skeletal muscle development, we first weighed the lean mass of both genotypes, and observed a significant decrease in *AKT2* KO mice. Further assessment showed decreased weight of hindlimb muscles, while the tibial length was equivalent in both genotypes, suggesting the decreased body mass is partly due to the loss of skeletal muscle despite WAT. Our further investigation of soleus muscle showed diminished cross-sectional area, and electromicroscopic images exhibited smaller sarcomeres of *AKT2* KO mice. Thus, our data suggests impaired skeletal muscle development due to *AKT2* deficiency.

To explore the mechanism involved in the regulation of skeletal development by AKT2, we focused on the MEF2 family, which is accepted as a key transcriptional factor during muscle differentiation and development (11, 50, 51). We observed that protein abundance of MEF2A was significantly suppressed in skeletal muscle of *AKT2* KO mice at both 3-month and 8-month old. Despite the important role of MEF2 family in skeletal muscle development, it is also known as a regulator of muscle energy metabolism (52, 53). It is reported that MEF2A can be activated by AMPK in skeletal muscle (37, 54). To explore if there is involvement of AMPK in AKT2-related MEF2A expression, AMPK activation was evaluated in skeletal muscle. Our results showed that p-AMPK was significantly decreased in skeletal muscle of *AKT2* KO mice either at the age of 3- or 8-months old. To verify the decreased AMPK phosphorylation is a direct effect of *AKT2* deficiency but not an indirect feedback, p-AMPK expression was checked in C2C12 cells transfecting with si*AKT2*. Our data showed that p-AMPK protein abundance decreased significantly when *AKT2* was knocked down in C2C12 cells, while AKT1 expression remained unchanged during the treatment. Therefore, our results suggest that AKT2 positively regulates AMPK activity at normal conditions in skeletal muscle. To further investigate the mechanisms underlying the role of AKT2 in skeletal muscle development, *AKT2* KO mice were treated with AICAR injection for 4 weeks. The result showed that after AICAR treatment, *AKT2* KO mice developed greatly alleviated insulin resistance; in addition, MEF2A was significantly induced in soleus muscle of *AKT2* KO mice. Thus, these results further confirmed that AKT2 regulates skeletal muscle development via AMPK activation.

Skeletal muscle, which accounts for 40% of body mass, is the major site of insulin-stimulated glucose uptake, with the majority of glucose that enters muscle fibers in response to insulin being stored as glycogen, and plays a key role in type 2 diabetes (55–57). Lack of *AKT2* in skeletal muscle decreased both the insulin sensitivity and responsiveness of glucose transport (15). Our results of *AKT2* KO and WT mice were in agreement with these reports. However, recently it has been reported that *AKT2* deficiency in skeletal muscle is not sufficient to affect development and glucose metabolism of skeletal muscle (58). One possible explanation for the discrepancy could be that, during the research, the authors used young mice at ages between 8-12 weeks. As we show in the present work, *AKT2* KO mice did not show significant decrease of body weight and glucose tolerance until 3-month of age. Therefore, we speculate that the lack of direct effects of AKT deletion in skeletal muscle reported in by *Jaiswal N et al.*, may be related to the age of the mice. Our study also proved that *AKT2* deficiency increased insulin resistance and hyperinsulinemia compared to WT during HFD-induced obesity, without obvious changes of islets. The protein expression of GLUT4 decreased in adipose tissue of obese and insulin resistant subjects (59, 60). However, the regulation of GLUT4 expression in skeletal muscle has not been well documented. Our data showed decreased GLUT4 expression in soleus muscle deficient for *AKT2* as well as in C2C12 cells with *AKT2* knockdown, indicating decreased glucose uptake into skeletal muscle cell, which may be a reason for impaired glucose tolerance and insulin resistance. In addition, there was no alteration of GLUT4 protein expression upon the stimulation of AMPK activator, suggesting a direct regulation of GLUT4 by AKT2. Therefore, these results suggest that AKT2 plays a key role in glucose homeostasis in skeletal muscle.

PPARα plays a central role in the regulation of muscle fuel preferences. Its expression also parallels the oxidative capacity of muscle fibers (61, 62). Therefore, our data confirmed the conclusion that *AKT2* deficiency impairs mitochondrial biogenesis, which in turn causes glycogen accumulation due to decreased glucose oxidation. Notably, there is research proved that *AKT2* deficiency did not cause alteration of p-AMPK or GLUT4 expression in muscle (63). However, the background of mice strain, also the gender and age of mice may cause the differences, as we showed that female mice exhibited delayed and alleviated glucose intolerance. Finally, to testify whether the phenomenon of sarcopenia and glucose metabolic disorder was caused via AKT2-AMPK signaling, by applying AICAR to *AKT2* KO mice, these mice exhibited ameliorated glucose tolerance and insulin resistance, and expression of proteins related to muscle development and mitochondrial biogenesis increased to normal level, suggesting AMPK is a key mediator during AKT2-associated regulation of skeletal muscle metabolism. In consistent with this result, we observed decreased total AKT2 protein abundance and increased p-AKT2 expression in soleus muscle of mice treated with HFD for 12 weeks, which exhibited glucose intolerance and insulin resistance. These data strengthened our conclusion that AKT2 and its activity plays a key role in glucose metabolism of skeletal muscle.

In summary, our study demonstrated an obligatory role of AKT2 in development and glucose metabolism of skeletal muscle. Our results showed that although *AKT2* deficiency is not lethal to life, however, its deficiency impairs systemic glucose metabolism. Moreover, AMPK phosphorylation promotes the transcription of molecules associated with skeletal muscle development and mitochondrial biogenesis. Our study gives new sights into the mechanism relating insulin resistance and sarcopenia of skeletal muscle in T2D, suggesting that AKT2 may be a key target for the therapy.

## Methods

### Animals

The *AKT2* knockout (*AKT2* KO) mouse line with C57/BL6 background was used in experiments. For all experiments, mice between 3 and 8 months old were used. The mice were maintained on a 12:12 h light/dark cycle, with controlled temperature (22 +/− 2 ℃) and humidity (55 +/− 5%) conditions. The mice received ad libitum access to water and standard lab chow, unless otherwise specified.

### Glucose tolerance test and insulin tolerance test

Glucose tolerance and insulin tolerance were investigated in *AKT2* KO and WT mice. Briefly, *AKT2* KO and WT mice were initially divided into a 3 and 8 months old group. For the glucose tolerance test (GTT), overnight-fasted male mice were intraperitoneally injected with glucose (2 g glucose/kg), and the glucose level was measured at 0、5、15、30、60、120 min. For the insulin tolerance test (ITT), insulin was intraperitoneally administered to male mice (fasting for 6 h) at 1 unit/kg, and glucose levels were monitored using blood collected from a tail vein with an Accu-Chek Performa glucometer (Roche, Basel, Switzerland). HOMA of insulin resistance (HOMA-IR) is calculated as follows: HOMA-IR=FINS/22.5e-InFPG (FINS: fasting serum insulin; FPG: fasting plasma glucose)

### Peritoneal injection of AICAR

6-month-old *AKT2* KO mice accepted peritoneal injection of AMPK activator AICAR (Toronto Research Chemicals Inc., North York, ON, Cananda, 0.5 mg/g of body weight) or saline once a day for continuous 4 weeks. Then the hearts were dissected for further investigation.

### Immunohistological staining

Skeletal muscle was routinely processed into paraffin sections. Sections of all cases were double-stained with haematoxylin and eosin stain (H&E) and periodic acid-Schiff stain (PAS). The paraffin section was stained in periodic acid solution for 15 minutes, washed with tap water and washed twice with distilled water. Then treated sliced with Schiff solution for 30 min, and rinsed for 5 min in running water. Subsequently, the section was put into hematoxylin for 1-3 min, then washed and differentiated. The sections were dehydrated and sealed. Microscopic examination, image acquisition and analysis. These sections were scanned by the digital slide scanner NanoZoomer 2.0RS (Hamamatsu), and the area of interstitial fibrosis was calculated by Image-Pro Plus 6.0 (Media Cybernetics, Rockville, Maryland, USA).

### Electron microscope

The embedded muscle tissue was sectioned and stained with uranium lead double staining (2% uranyl acetate saturated alcohol solution, lead citrate, each stained for 15 min), and the sections were dried overnight at room temperature. Observed under a transmission electron microscope (Hitachi), image analysis was acquired.

### Glycogen content in muscle tissue

Glycogen in skeletal muscle was measured by a glycogen content analysis kit (QIYI Biological Technology, ShangHai, China, QYS-234078). About 50 mg of minced skeletal muscle samples were incubated with concentrated alkali in a test tube. The samples were minced and homogenized with a homogenizer. Homogenates were heated in 95℃ water bath for 20 min. Then the samples were diluted with distilled water after cooling. Homogenates were centrifuged by 8000 g for 10 min at 25℃. The supernatant was mixed with concentrated sulfuric acid, placed in 95°C water bath, then cooled with cool water. Mixture was transferred to a 96-well plate, and the absorbance was measured at a wavelength of 620 nm. Glycogen concentration was determined by referring a glycogen standard solution in the kit.

### Quantification of mitochondrial DNA (mtDNA) copy number

Total DNA extraction from isolated skeletal muscle tissue was carried out using the genomic DNA extraction Kit (Beyotime, D0063) according to the manufacturer’s recommendations. To determine the mtDNA copy number in skeletal muscle tissue, real-time quantitative PCR (Q-PCR) was performed using the Stepone System in a 10 µl reaction volume containing 5 μL Master Mix,0.3 μL forward primer (10 pMol), 0.3 μL reverse primer (10 pMol), 3.9 μL nuclease-free H2O, and 0.5 μL genomic DNA. The reactions were performed as follows: initial denaturing at 95°C for 10 min, and 40 cycles at 95℃ for 15 min, 60℃ for 30 s and proceeded by an initial denaturation of 10 min at 95℃ and followed by a continuous melt curve from 60 to 95℃. A melting curve was analyzed in order to check the specificity of the PCR product. Mitochondrial gene *nd-5* was analyzed to assess mtDNA copy number. All samples were amplified in triplicate. Results were expressed as ratio of *nd-5* to *β-actin*.

### C2C12 myoblasts culture, differentiation and Transfection

C2C12 cells were cultured in growth medium (GM) consisting of Dulbecco’s modified Eagle’s medium (DMEM, sigma, USA) with 10% foetal bovine serum, 100 U/mL penicillin and 100 µg/mL streptomycin under a humidified atmosphere of 5% CO2 at 37°C. To induce differentiation, the cells were grown to 70-80% confluent, and the medium was then replaced with differentiation medium (DM) consisting of DMEM containing 2% horse serum (Gibco, USA), 100 U/mL penicillin and 100 µg/mL streptomycin, which was replenished everyday. After 3 days, they were transfected with mouse AKT2-targeting siRNA (si*AKT2*) or siRNA for negative control (NC). On the fourth day after transfection, cells were collected for protein extraction.

### Western Blot

To prepare protein samples from the cultured cells, PBS-washed cells were lysed in lysis buffer (125mM Tris, 2% SDS at pH 6.8). Mouse tissue samples were snap-frozen in liquid nitrogen and maintained at –80°C. Tissue lysates were prepared in lysis buffer (125 mmol/L Tris, 2% SDS, pH6.8). Lysis buffer supplemented with a proteinase inhibitor (Roche, Basel, Switzerland) and phosphatase inhibitor cocktail I (MCE, HY-K0021) and phosphatase inhibitor cocktail II (MCE, HY-K0022). Approximately 40 mg of protein sample was separated via 8% SDS–PAGE and then transferred onto polyvinylidene difluoride membrane (Millipore, Bedford, Massachusetts, USA). Then the membranes were blocked with 2% BSA (bovine serum albumin) in TBST (50 mmol/L Tris, 150 mmol/L NaCl, 0.5 mmol/L tris-buffered saline and Tween-20, pH 7.5). After blocking, the blots were incubated with antibodies (Table 1) overnight at 4℃. Following, samples where incubated with appropriate HPR conjugated secondary antibodies. The protein bands were detected with enhanced chemiluminescence kit (Thermo Scientific, USA). The images were quantitatively analyzed by using the Image J program to make comparisons between different groups.

**Table 1.**
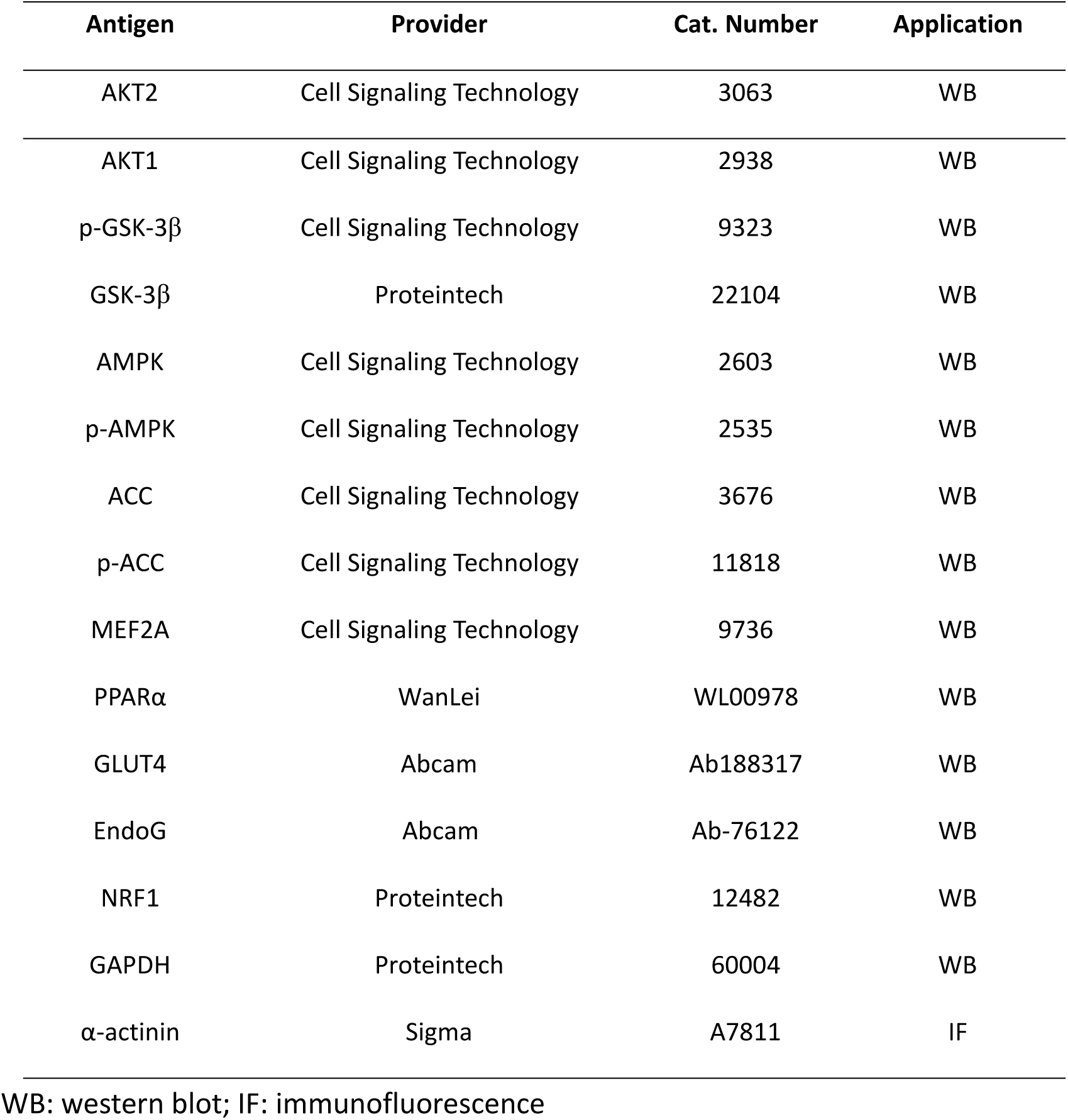
Antibodies used for western blot and IF

### Total RNA extraction and and reverse transcription-polymerase chain reaction

Total RNA was extracted using the TRIeasy reagent (Yeasen, 10606ES60) according to the normal instructions. A 1 μg sample of total RNA was reverse-transcribed using a reverse transcription system kit (abm, G486) according to the operational instruction. Then real-time PCR was performed with cDNA templates and SYBR Green qPCR Master Mix (Selleck, B21403) by using StepOnePlus Real-Time PCR (Applied Biosystems, Foster City, California, USA). Briefly, sequences were amplified using 5ul Master Mix, 0.3 μl forward primer (10 pmol) (Table 2) of *pgc-1α*, *nrf-1*, *pparα*, *mtfa*, *gapdh*. 3.9 μl nuclease-free H2O and 0.5 μl cDNA in a total volume of 10 μl. PCR conditions were 40 cycles of 15 sat 95℃, 30 s at 60 ℃, proceeded by an initial denaturation of 10 min at 95℃ and followed by a continuous melt curve from 60 to 95℃. All samples were amplified in triplicate, and the results were normalized to the expression of 18s rRNA.

**Table 2.**
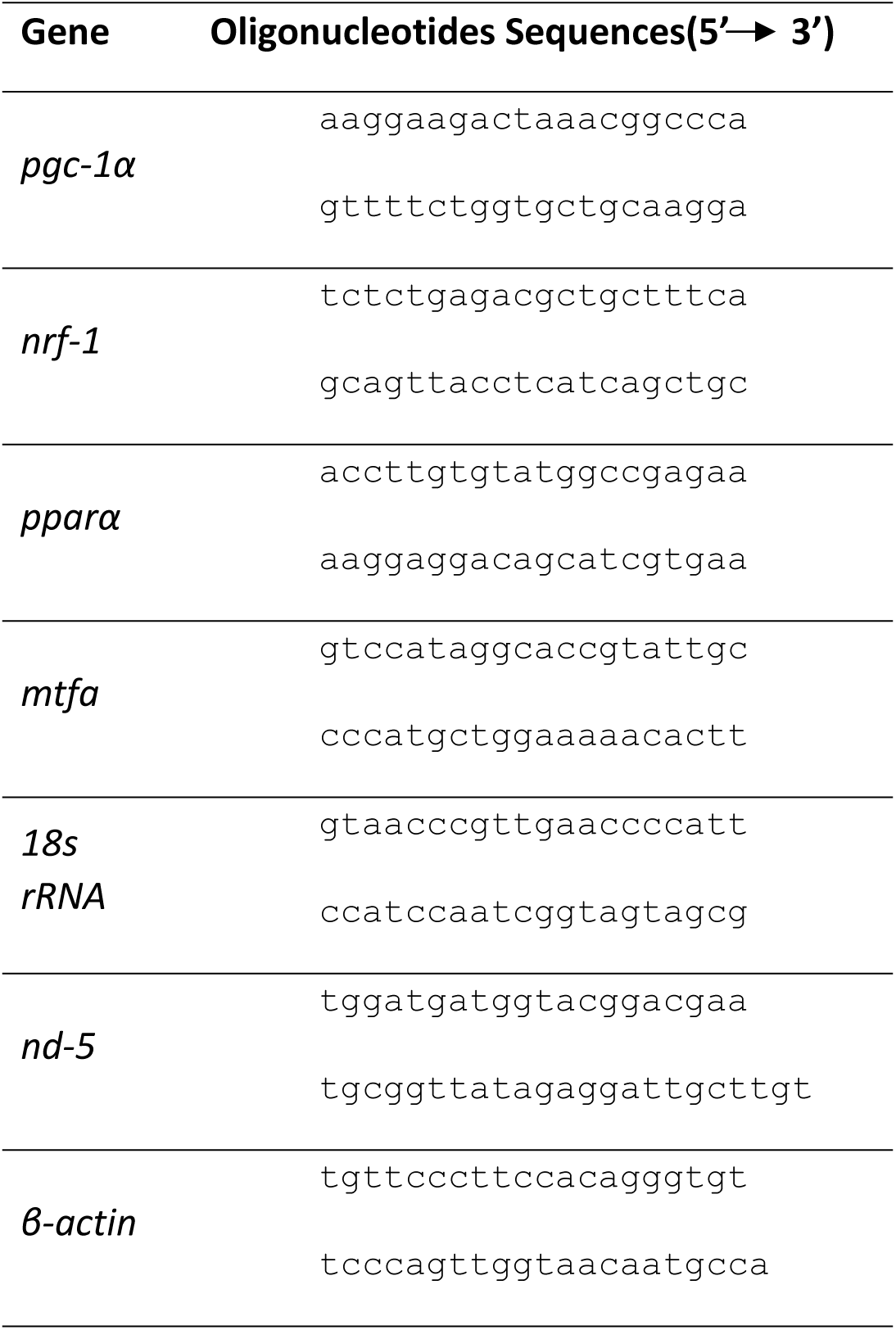
Primers used for real-time PCR

### Immunofluorescence

C2C12 cells replaced with differentiation medium (DM) for 3 days, and then were transfected with mouse si*AKT2* or NC. On the fourth day after transduction of lentivirus, cells were used for immunofluorescence detection. C2C12 cells were grown in a 4-well plate (NUNC, Roskilde, Denmark) and fixed with 4% paraformaldehyde, rinsed twice with PBS. Then C2C12 cell was incubated in blocking solution (BS, 2% horse serum, 0.1% Triton, 2% BSA) for 1 h at room temperature (RT). Primary antibodies (α-actinin) were diluted in BS and incubated for 2 h at room temperature. Then the wells were washed three times with BS and incubated with the secondary antibody for 1 h at RT. Finally, cell nuclei were stained with Hoechst 33342 (SIGMA-ALDRICH, #B2883) for 30 min at room temperature. Cells were rinsed twice with PBS and mounted in Vectashield (Vecto Laboratories, Burlingame, CA, USA), which then were observed with inverted fluorescence microscope (Carl ZEISS, Axio Vert. A1, Dublin, CA, USA).

### Statistics

Data are shown as mean ± SEM (standard error of mean). All statistical analyses were performed using Prism (Graphpad, San Diego, CA, USA) software. Differences between two groups were compared by unpaired 2-tailed Student’s *t* test. Multiple groups were tested via two-way ANOVA followed by Bonferroni post-test. A *p*-value < 0.05 was considered statistically significant.

## Author contributions

Junmei Ye, Fangrong Yan and Daniel Sanchis conceived and designed the experiments. Miao Chen, Caoyu Ji, Fei Xiao, Dandan Chen, Shuya Gao, Qingchen Yang, Yue Peng collected, analyzed, and interpreted data. Junmei Ye drafted the article, Daniel Sanchis and Fangrong Yan revised the manuscript.

## Acknowledgments

This work was supported by the Natural Science Foundation of Jiangsu Province (Grant No. BK20191324); “Double First-Class” University project (Grant No. CPU2018GY09); National Natural Science Foundation of China (Grant No. 81973145).

## References

1. BӧHni R, ., Riesgo-Escovar J, ., Oldham S, ., Brogiolo W, ., Stocker H, ., Andruss BF, et al. Autonomous control of cell and organ size by CHICO, a Drosophila homolog of vertebrate IRS1-4. Cell. 1999;97(7):865–75.

2. Mccurdy CE, and Cartee GD. Akt2 is essential for the full effect of calorie restriction on insulin-stimulated glucose uptake in skeletal muscle. Diabetes. 2005;54(5):1349–56.

3. Brazil DP, and Hemmings BA. Ten years of protein kinase B signalling: a hard Akt to follow. Trends in Biochemical Sciences. 2001;26(11):657–64.

4. Coffer P, Jin J, and Woodgett J. Protein kinase B (c-Akt): a multifunctional mediator of phosphatidylinositol 3-kinase activation. Biochemical Journal. 1998;335(1):1–13.

5. Cho H, Thorvaldsen JL, Chu Q, Feng F, and Birnbaum MJ. Akt1/PKB? Is Required for Normal Growth but Dispensable for Maintenance of Glucose Homeostasis in Mice. Journal of Biological Chemistry.

6. Cho H, Mu J, Kim JK, Thorvaldsen JL, Chu Q, Crenshaw EB, 3rd, et al. Insulin resistance and a diabetes mellitus-like syndrome in mice lacking the protein kinase Akt2 (PKB beta). Science. 2001;292(5522):1728–31.

7. Easton RM, Han C, Kristin R, Shineman DW, Moshe M, Forman MS, et al. Role for Akt3/protein kinase Bgamma in attainment of normal brain size. Molecular & Cellular Biology. 2005;25(5):1869–78.

8. Wan M, Leavens K, Saleh D, Easton R, Guertin D, Peterson T, et al. Postprandial Hepatic Lipid Metabolism Requires Signaling through Akt2 Independent of the Transcription Factors FoxA2, FoxO1, and SREBP1c. Cell Metabolism. 2011;14(4):516–27.

9. Chen D, Chen F, Xu Y, Zhang Y, Li Z, Zhang H, et al. AKT2 deficiency induces retardation of myocyte development through EndoG-MEF2A signaling in mouse heart. Biochem Biophys Res Commun. 2017;493(4):1410–7.

10. Montserrat N, ., Sánchez-Gurmaches J, ., Serrana D, García De La, Navarro MI, and Gutiérrez J,. IGF-I binding and receptor signal transduction in primary cell culture of muscle cells of gilthead sea bream: changes throughout in vitro development. Cell & Tissue Research. 2007;330(3):503–13.

11. Ji-Hoon L, Rhonda BD, and Olson EN. Heart- and muscle-derived signaling system dependent on MED13 and Wingless controls obesity in Drosophila. Proceedings of the National Academy of Sciences of the United States of America. 2014;111(26):9491–6.

12. Mora S, ., and Pessin JE. The MEF2A isoform is required for striated muscle-specific expression of the insulin-responsive GLUT4 glucose transporter. Journal of Biological Chemistry. 2000;275(21):16323–8.

13. Potthoff MJ, Arnold MA, John MA, Richardson JA, Rhonda BD, and Olson EN. Regulation of skeletal muscle sarcomere integrity and postnatal muscle function by Mef2c. Molecular & Cellular Biology. 2007;27(23):8143–51.

14. Potthoff MJ, Hai W, Arnold MA, Shelton JM, Johannes B, John MA, et al. Histone deacetylase degradation and MEF2 activation promote the formation of slow-twitch myofibers. Journal of Clinical Investigation. 2007;117(9):2459–67.

15. Garofalo RS, Orena SJ, Kristina R, Torchia AJ, Stock JL, Hildebrandt AL, et al. Severe diabetes, age-dependent loss of adipose tissue, and mild growth deficiency in mice lacking Akt2/PKB beta. Journal of Clinical Investigation. 2003;112(2):197–208.

16. Mccurdy CE, Simon S, Holliday MJ, Andrew P, Houck JA, David P, et al. Attenuated Pik3r1 expression prevents insulin resistance and adipose tissue macrophage accumulation in diet-induced obese mice. Diabetes. 2012;61(10):2495–505.

17. Caroline T, Samira A, Rim H, Yun X, Birnbaum MJ, and Mario P. The combined deletion of S6K1 and Akt2 deteriorates glycemic control in a high-fat diet. Molecular & Cellular Biology. 2012;32(19):4001–11.

18. Kuninger D, Wright A, and Rotwein P. Muscle cell survival mediated by the transcriptional coactivators p300 and PCAF displays different requirements for acetyltransferase activity. American Journal of Physiology. 2006;291(4):699–709.

19. Sato S, Ogura Y, Tajrishi MM, and Kumar A. Elevated levels of TWEAK in skeletal muscle promote visceral obesity, insulin resistance, and metabolic dysfunction. FASEB J. 2015;29(3):988–1002.

20. Defronzo RA. From the Triumvirate to the Ominous Octet: A New Paradigm for the Treatment of Type 2 Diabetes Mellitus. Diabetes. 2009.

21. Gosmanov AR, Zheng F, Xianqiang M, Schneider EG, and Thomason DB. ATP-sensitive potassium channels mediate hyperosmotic stimulation of NKCC in slow-twitch muscle. Am J Physiol Cell Physiol. 2004;286(3):586–95.

22. Brozinick JT, Roberts BR, and G Lynis D. Defective signaling through Akt-2 and -3 but not Akt-1 in insulin-resistant human skeletal muscle: potential role in insulin resistance. Diabetes. 2003;52(4):935.

23. Gowans GJ, and D Grahame H. AMPK: a cellular energy sensor primarily regulated by AMP. Biochem Soc Trans. 2014;42(1):71–5.

24. Russell RR, Bergeron R, ., Shulman GI, and Young LH. Translocation of myocardial GLUT-4 and increased glucose uptake through activation of AMPK by AICAR. American Journal of Physiology. 1999;277(2 Pt 2):H643.

25. J?Rgensen SB, Nielsen JN, Birk JB, Grith Skytte O, Benoit V, Fabrizio A, et al. The alpha2-5’AMP-activated protein kinase is a site 2 glycogen synthase kinase in skeletal muscle and is responsive to glucose loading. Diabetes. 2004;53(12):3074–81.

26. Bolster DR, Crozier SJ, Kimball SR, and Jefferson LS. AMP-activated protein kinase suppresses protein synthesis in rat skeletal muscle through down-regulated mammalian target of rapamycin (mTOR) signaling. Journal of Biological Chemistry. 2002;277(27):23977–80.

27. Vladimir L, Pedro M, Matthew B, Louise B, Shiemaa K, Lunde JA, et al. Chronic AMPK activation evokes the slow, oxidative myogenic program and triggers beneficial adaptations in mdx mouse skeletal muscle. Human Molecular Genetics. 2011;20(17):3478–93.

28. Bergeron R, ., Ren JM, Cadman KS, Moore IK, Perret P, ., Pypaert M, ., et al. Chronic activation of AMP kinase results in NRF-1 activation and mitochondrial biogenesis. Am J Physiol Endocrinol Metab. 2001;281(6):E1340.

29. Winder WW, Holmes BF, Rubink DS, Jensen EB, Chen M, and Holloszy JO. Activation of AMP-activated protein kinase increases mitochondrial enzymes in skeletal muscle. J Appl Physiol (1985). 2000;88(6):2219–26.

30. Haihong Z, Ming RJ, Young LH, Marc P, James M, Birnbaum MJ, et al. AMP kinase is required for mitochondrial biogenesis in skeletal muscle in response to chronic energy deprivation. Proceedings of the National Academy of Sciences of the United States of America. 2002;99(25):15983–7.

31. Suzanne K, Soltys CLM, Barr AJ, Ichiro S, Kenneth W, and Dyck JRB. Akt activity negatively regulates phosphorylation of AMP-activated protein kinase in the heart. Journal of Biological Chemistry. 2003;278(41):39422–7.

32. Annett HW, Veronique N, Chia-Chen C, Skeen JE, Nahum S, and Nissim H. Akt activates the mammalian target of rapamycin by regulating cellular ATP level and AMPK activity. Journal of Biological Chemistry. 2005;280(37):32081–9.

33. Holmes BF, Sparling DP, Olson AL, Winder WW, and Dohm GL. Regulation of muscle GLUT4 enhancer factor and myocyte enhancer factor 2 by AMP-activated protein kinase. Am J Physiol Endocrinol Metab. 2005;289(6):E1071–6.

34. Goncalves MD, Pistilli EE, Anthony B, Birnbaum MJ, Jennifer L, Khurana TS, et al. Akt deficiency attenuates muscle size and function but not the response to ActRIIB inhibition. Plos One. 2010;5(9):e12707.

35. Ren J, Yang L, Zhu L, Xu X, Ceylan AF, Guo W, et al. Akt2 ablation prolongs life span and improves myocardial contractile function with adaptive cardiac remodeling: role of Sirt1-mediated autophagy regulation. Aging Cell. 2017;16(5):976.

36. Leavens KF, Easton RM, Shulman GI, Previs SF, and Birnbaum MJ. Akt2 is required for hepatic lipid accumulation in models of insulin resistance. Cell Metabolism. 2009;10(5):405–18.

37. Gan Z, Burkart-Hartman EM, Han DH, Finck B, Leone TC, Smith EY, et al. The nuclear receptor PPARbeta/delta programs muscle glucose metabolism in cooperation with AMPK and MEF2. Genes Dev. 2011;25(24):2619–30.

38. Chris MDR, Junmei Y, Rizwan A, Xi-Ming S, Anna S, James W, et al. Endonuclease G is a novel determinant of cardiac hypertrophy and mitochondrial function. Nature. 2011;478(7367):114.

39. Koh JH, Hancock CR, Terada S, Higashida K, Holloszy JO, and Han DH. PPARbeta Is Essential for Maintaining Normal Levels of PGC-1alpha and Mitochondria and for the Increase in Muscle Mitochondria Induced by Exercise. Cell Metab. 2017;25(5):1176–85 e5.

40. Wu Z, Puigserver P, Andersson U, Zhang C, Adelmant G, Mootha V, et al. Mechanisms controlling mitochondrial biogenesis and respiration through the thermogenic coactivator PGC-1. Cell. 1999;98(1):115–24.

41. Puigserver P, Wu Z, Park CW, Graves R, Wright M, and Spiegelman BM. A cold-inducible coactivator of nuclear receptors linked to adaptive thermogenesis. Cell. 1998;92(6):829–39.

42. Lezza AM, Pesce V, Cormio A, Fracasso F, Vecchiet J, Felzani G, et al. Increased expression of mitochondrial transcription factor A and nuclear respiratory factor-1 in skeletal muscle from aged human subjects. FEBS Lett. 2001;501(1):74–8.

43. Thiebaud D, ., Jacot E, ., Defronzo RA, Maeder E, ., Jequier E, ., and Felber JP. The effect of graded doses of insulin on total glucose uptake, glucose oxidation, and glucose storage in man. Diabetes. 1982;31(11):957–63.

44. Klip A, and Bonen A. The many ways to regulate glucose transporter 4. Applied Physiology Nutrition & Metabolism. 2009;34(3):481–7.

45. Yaluri N, Modi S, and Kokkola T. Simvastatin induces insulin resistance in L6 skeletal muscle myotubes by suppressing insulin signaling, GLUT4 expression and GSK-3β phosphorylation. Biochemical & Biophysical Research Communications. 2016;480(2):194–200.

46. Kei S, Arnolds DE, Nobuharu F, Kramer HF, Hirshman MF, and Goodyear LJ. Role of Akt2 in contraction-stimulated cell signaling and glucose uptake in skeletal muscle. Am J Physiol Endocrinol Metab. 2006;291(5):1031–7.

47. Muslin AJ. Akt2: a critical regulator of cardiomyocyte survival and metabolism. Pediatr Cardiol. 2011;32(3):317–22.

48. Etzion S, Etzion Y, DeBosch B, Crawford PA, and Muslin AJ. Akt2 deficiency promotes cardiac induction of Rab4a and myocardial beta-adrenergic hypersensitivity. J Mol Cell Cardiol. 2010;49(6):931–40.

49. Yang JY, Deng W, Chen Y, Fan W, Baldwin KM, Jope RS, et al. Impaired translocation and activation of mitochondrial Akt1 mitigated mitochondrial oxidative phosphorylation Complex V activity in diabetic myocardium. J Mol Cell Cardiol. 2013;59:167–75.

50. Taylor MV. Skeletal muscle development on the 30th Anniversary of MyoD. Seminars in Cell & Developmental Biology. 2017;72:1–2.

51. Kim MS, Fielitz J, McAnally J, Shelton JM, Lemon DD, McKinsey TA, et al. Protein kinase D1 stimulates MEF2 activity in skeletal muscle and enhances muscle performance. Mol Cell Biol. 2008;28(11):3600–9.

52. Michael LF, Wu Z, ., Cheatham RB, Puigserver P, ., Adelmant G, ., Lehman JJ, et al. Restoration of insulin-sensitive glucose transporter (GLUT4) gene expression in muscle cells by the transcriptional coactivator PGC-1. Proceedings of the National Academy of Sciences of the United States of America. 2001;98(7):3820–5.

53. Naya FJ, Black BL, Wu H, Bassel-Duby R, Richardson JA, Hill JA, et al. Mitochondrial deficiency and cardiac sudden death in mice lacking the MEF2A transcription factor. Nature Medicine. 2002;8(11):1303–9.

54. Holmes BF, Sparling DP, Olson AL, Winder WW, and Dohm GL. Regulation of muscle GLUT4 enhancer factor and myocyte enhancer factor 2 by AMP-activated protein kinase. Am J Physiol Endocrinol Metab. 2005;289(6):E1071.

55. Bouskila M, Hirshman MJ, and Goodyear LJ. Insulin promotes glycogen synthesis in the absence of GSK3 phosphorylation in skeletal muscle. Am J Physiol Endocrinol Metab. 2008;294(1):E28.

56. Shulman GI, Rothman DL, Jue T, ., Stein P, ., Defronzo RA, and Shulman RG. Quantitation of muscle glycogen synthesis in normal subjects and subjects with non-insulin-dependent diabetes by 13C nuclear magnetic resonance spectroscopy. N Engl J Med. 1990;322(4):223–8.

57. Fu X, Zhu M, Zhang S, Foretz M, Viollet B, and Du M. Obesity Impairs Skeletal Muscle Regeneration Through Inhibition of AMPK. Diabetes. 2016;65(1):188–200.

58. Jaiswal N, Gavin MG, Quinn WJ, 3rd, Luongo TS, Gelfer RG, Baur JA, et al. The role of skeletal muscle Akt in the regulation of muscle mass and glucose homeostasis. Mol Metab. 2019.

59. Kelly KR, Kashyap SR, O’Leary VB, Major J, Schauer PR, and Kirwan JP. Retinol-binding protein 4 (RBP4) protein expression is increased in omental adipose tissue of severely obese patients. Obesity. 2012;18(4):663–6.

60. Hussey SE, Mcgee SL, Garnham A, ., Wentworth JM, Jeukendrup AE, and Hargreaves M,. Exercise training increases adipose tissue GLUT4 expression in patients with type 2 diabetes. Diabetes Obesity & Metabolism. 2011;13(10):959–62.

61. Finck BN, Bernal-Mizrachi C, Dong HH, Coleman T, Sambandam N, Lariviere LL, et al. A potential link between muscle peroxisome proliferator- activated receptor-α signaling and obesity-related diabetes. Cell Metabolism. 2005;1(2):133–44.

62. Muoio DM, Way JM, Tanner CJ, Winegar DA, Kliewer SA, Houmard JA, et al. Peroxisome proliferator-activated receptor-alpha regulates fatty acid utilization in primary human skeletal muscle cells. Diabetes. 2002;51(4):901–9.

63. Jain SS, Snook LA, Glatz JFC, Luiken JJFP, Holloway GP, Thurmond DC, et al. Munc18c provides stimulus-selective regulation of GLUT4 but not fatty acid transporter trafficking in skeletal muscle. Febs Letters. 2012;586(16):2428–35.

