## Supplementary material for "AKT2 deficiency causes sarcopenia and metabolic disorder of skeletal muscle": Supplimental Figures

Suppl. Figure 1

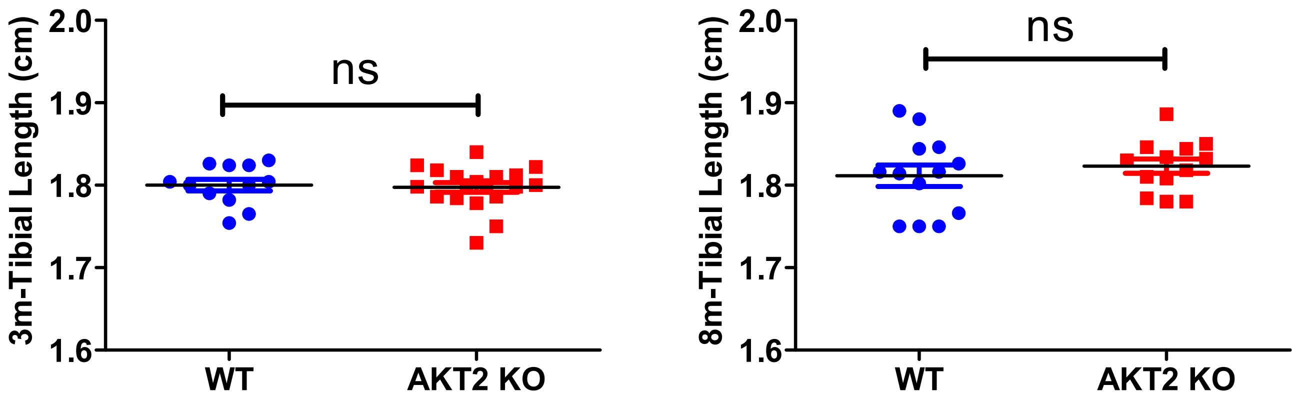


**Suppl. Figure 1. Tibial length of WT and AKT2 KO mice.** Tibia was dissected from mice of both genotypes at the age of 3-month-old (left panel) (WT, n=12; AKT2 KO, n=19) and 8-month-old (right panel) (WT, n=13; AKT2 KO, n=13) was assessed. ns, not significant, WT versus KO. Data are presented as mean ± SEM.

Suppl. Figure 2

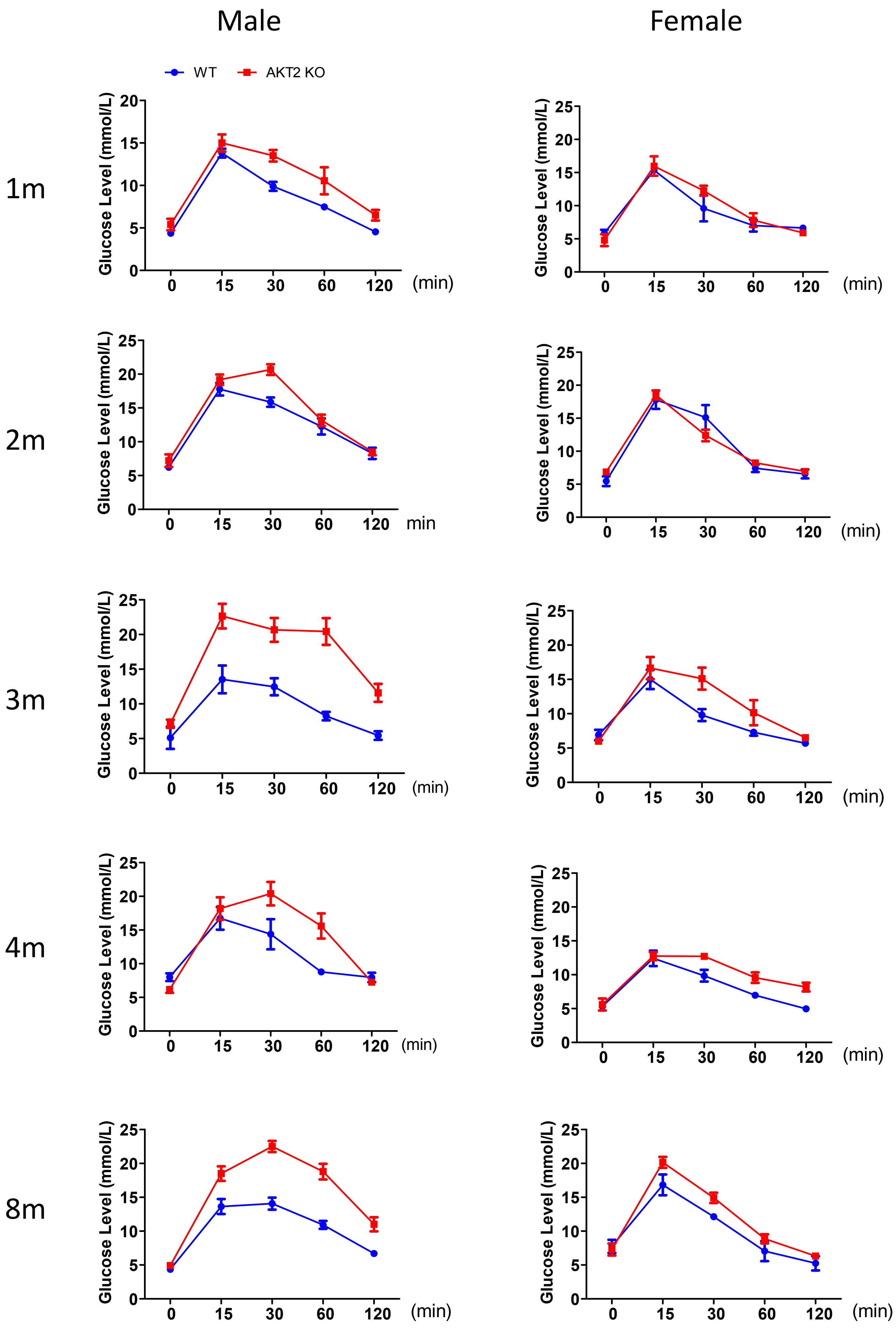


**Suppl. Figure 2. Glucose tolerance of WT and AKT2 KO mice.** IP-GTT was performed to assess glucose tolerance of males and females of both WT and AKT2 KO mice at different ages that ranges from 1-month-old to 8-month-old (n=5 of males and females for both genotypes).

Suppl. Figure 3

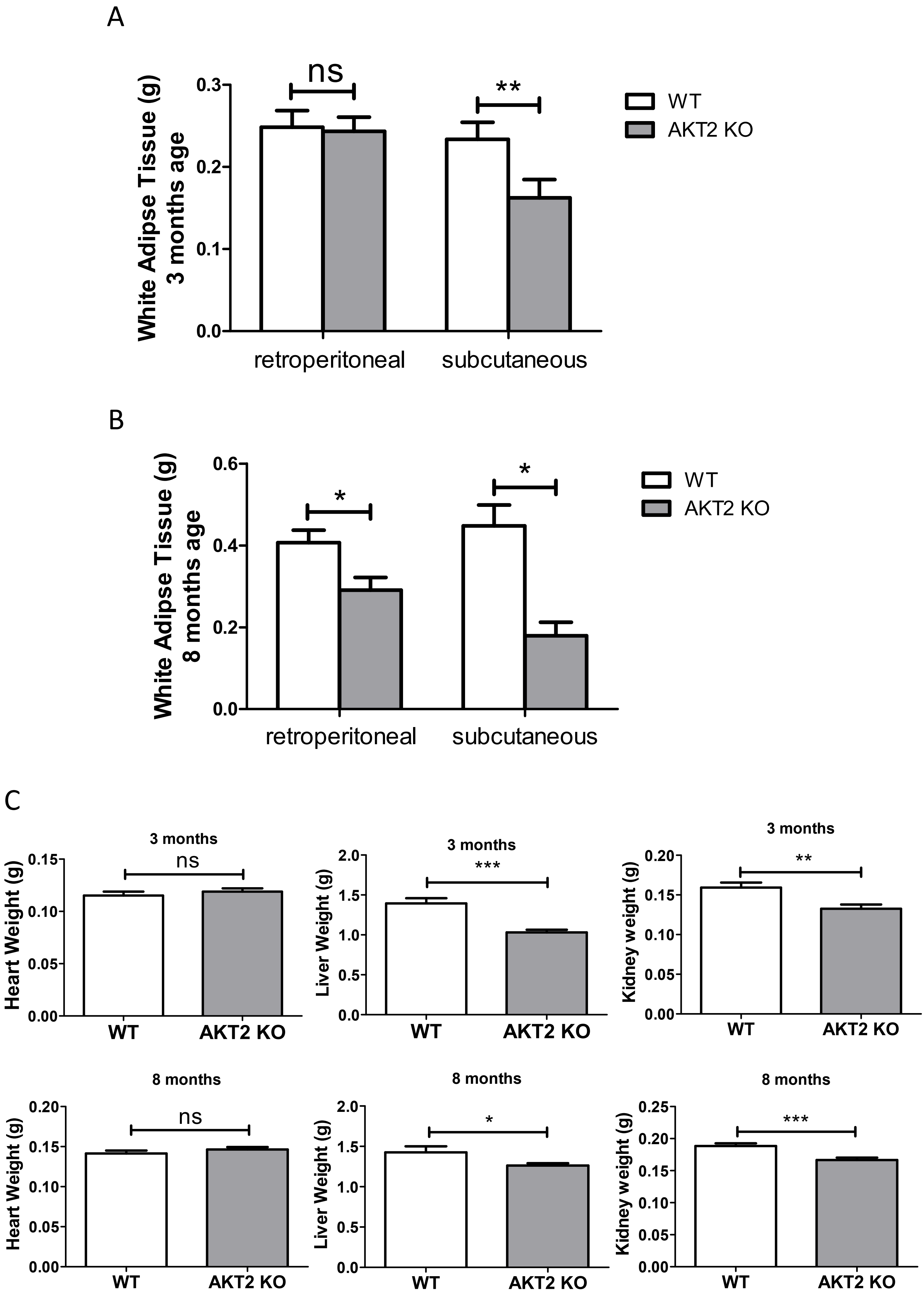


**Suppl. Figure 3. Effect of AKT2 deficiency on visceral mass.** (A and B) Retroperitoneal WAT and subcutaneous WAT mass of WT and AKT2 KO mice at the age of 3- (WT, n=6; AKT2 KO, n=11) and 8-month-old (n=10 for both genotypes). (C) Weight of heart, liver and kidney from WT and AKT2 KO mice at the age of 3-month-old (upper panel) (WT, n=6; AKT2 KO, n=11) and 8-month-old (lower panel) (n=10 for both genotypes). *P < 0.05; **P < 0.01; ***P < 0.001; ns, not significant, WT versus KO. Data are presented as mean ± SEM.

Suppl. Figure 4

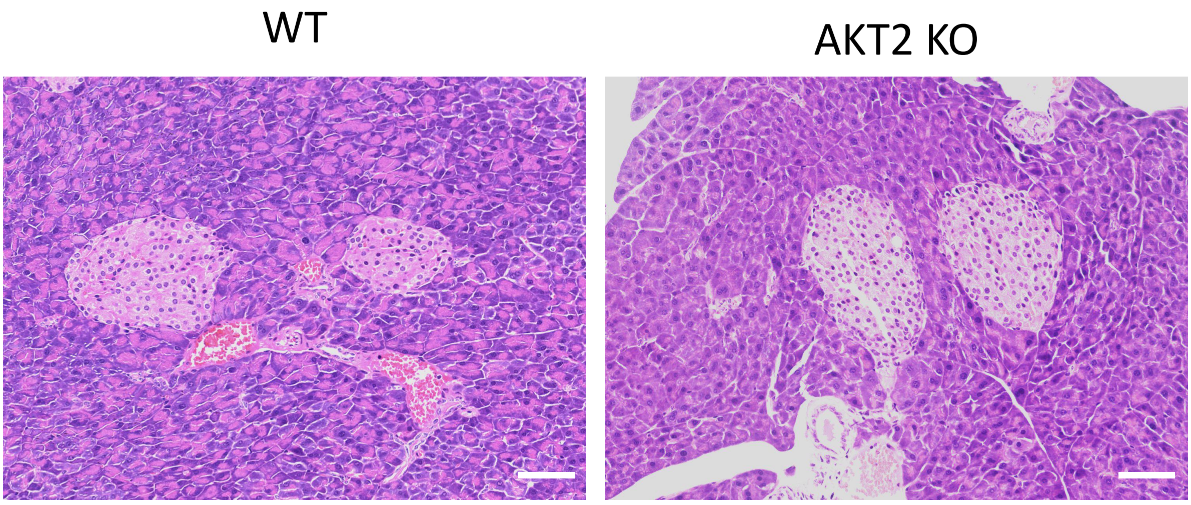


**Suppl. Figure 4. Effect of AKT2 deficiency on morphology of islets.** Representative images of pancreas H & E staining showing no obvious changes of islet in AKT2 KO mice compared to that of WT (n=4 per genotype). scale bar: 100 μm.

Suppl. Figure 5

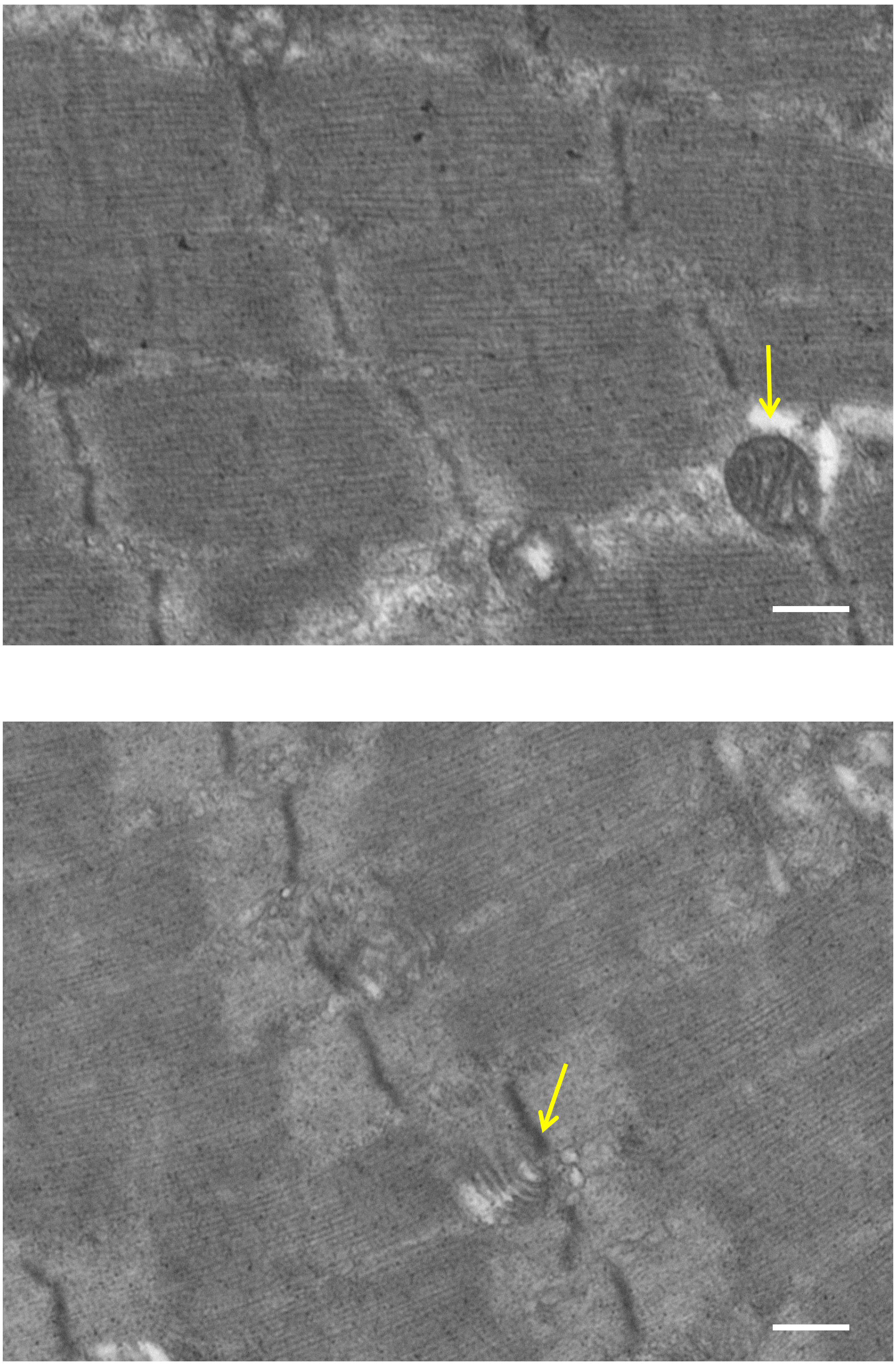


**Suppl. Figure 5. Sarcomeric array and mitochondria morphology.** Representative transmission electron microscopic images showing regular sarcomeric array and mitochondrial morphology in WT and AKT2 KO soleus muscle (n=3 per genotype). scale bar: 4 µm.
